# Expression Atlas of Avian Neural Crest Proteins: Neurulation to Migration

**DOI:** 10.1101/2021.08.17.456488

**Authors:** Brigette Y. Monroy, Carly J. Adamson, Alexis Camacho-Avila, Christian N. Guerzon, Camilo V. Echeverria, Crystal D. Rogers

## Abstract

Neural crest (NC) cells are a dynamic population of embryonic stem cells that create various adult tissues in vertebrate species including craniofacial bone and cartilage and the peripheral and enteric nervous systems. NC development is a conserved and complex process that is controlled by a tightly regulated gene regulatory network (GRN) of morphogens, transcription factors, and cell adhesion proteins. While multiple studies have characterized the expression of several GRN factors in single species, a comprehensive protein analysis that directly compares expression across development is lacking. To address this, we used three closely related avian models, *Gallus gallus* (chicken), *Coturnix japonica* (Japanese quail), and *Pavo cristatus* (Indian peafowl), to compare the localization and timing of four GRN transcription factors, PAX7, SOX9, SNAI2, and SOX10 from the onset of neurulation to migration. While the spatial expression of these factors is largely conserved, we find that quail NC cells express SOX9, SNAI2, and SOX10 proteins at the equivalent of earlier developmental stages than chick and peafowl. In addition, quail NC cells migrate farther and more rapidly than the larger organisms. These data suggest that despite a conservation of NC GRN players, differences in the timing of NC development between species remain a significant frontier to be explored with functional studies.

**Graphical abstract:** Comparative analysis of neural crest (NC) protein spatiotemporal localization in quail, chick, and peafowl embryos.
Avian embryos were incubated for different lengths of time to achieve the same developmental stage marked by somite number (somite stage, SS) as described by Hamburger and Hamilton (HH) in 1951. (A) Quail, (B) chick, and (C) peafowl embryos were collected for immunohistochemistry (IHC) to define and quantify the timeline of NC protein expression. We specifically focused on the expression of PAX7, SNAI2, SOX9, and SOX10 proteins. We determined that neural crest development as marked by common NC-specific proteins differs between species. Rather than similar, but scaled development, each organism has its own NC developmental timeline. Quail embryos develop much more rapidly than their counterparts, chick and peafowl. Further, chick and peafowl amino acid sequences are more similar to each other than they are to quail.

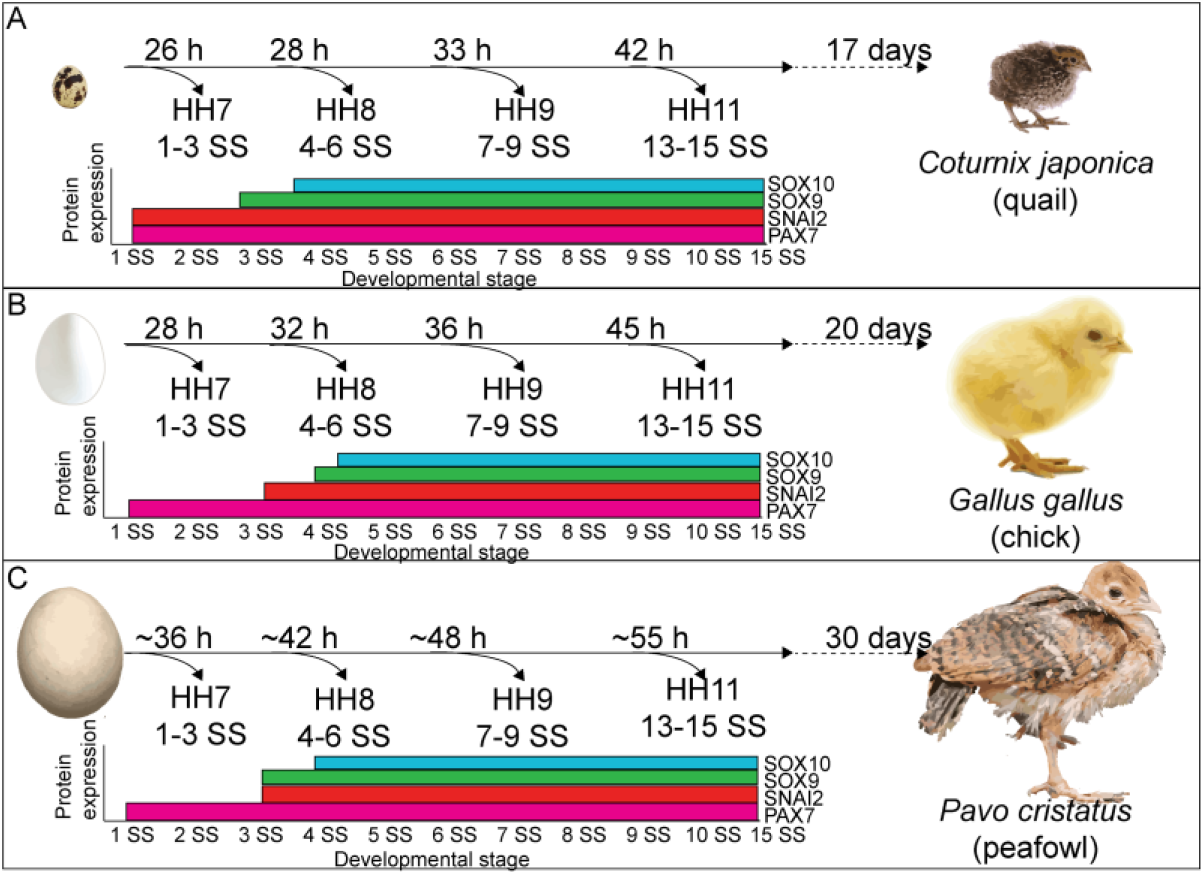

## 1. INTRODUCTION

Neural crest (NC) cells are a multipotent population of cells that arise from the epithelial ectodermal germ layer, undergo an epithelial to mesenchymal transition (EMT), migrate, and differentiate into many different cell and tissue derivatives (Martik and Bronner, 2017). During early neurulation, the lateral edges of the elevated neural plate border begin to express NC progenitor proteins, including PAX7 (Basch et al., 2006). Then, as the neural plate border fuses together to form the neural tube, NC cells are specified in the most dorsal region and begin to upregulate definitive NC markers, such as SNAI2 and SOX9 (Cheung and Briscoe, 2003; Spokony et al., 2002; Taneyhill et al., 2007). As development progresses, avian NC cells activate SOX10 prior to delaminating from the neural tube and initiate an EMT driven by changes in the expression of transcriptional regulators of cell adhesion molecules, cell adhesion, and in cell polarity (Cheng et al., 2000; Honore et al., 2003). During EMT and early migration, premigratory NC cells express E-cadherin (Dady et al., 2012; Dady and Duband, 2017; Rogers et al., 2018) and upregulate the expression of migratory cadherins (Cadherin-11 and Cadherin-7) (Kawano et al., 2002; Manohar et al., 2020; Vallin et al., 1998). Following EMT, migratory NC cells reduce expression of E-Cadherin, modulating collective to mesenchymal migration downstream of SIP1/ZEB2 (Rogers et al., 2013) and acquire a mesenchymal and migratory state. Next, they travel through distinct pathways to reach a variety of destinations, where they undergo differentiation (Bronner and LeDouarin, 2012). Any abnormalities that arise during the development of NC cells can lead to a variety of craniofacial birth defects, as well as a number of rare neurocristopathies (Pilon, 2021).

NC cell development is regulated by a complex gene regulatory network (GRN) consisting of secreted factors that drive rapid changes in the expression of transcription factors, which then regulate the expression of cadherins (Simoes-Costa and Bronner, 2015). While there are hundreds of genes with reported functions in the GRN, there is still a lack of information about the endogenous spatiotemporal localization of their encoded proteins across species. Here, we quantify the onset, localization, and timing of NC cell development via comparative analysis of GRN proteins in three avian species, *Gallus gallus* (chicken), *Coturnix japonica* (quail), and *Pavo cristatus* (peafowl) by focusing on the expression of PAX7, SOX9, SNAI2, and SOX10. PAX7, a paired-box transcription factor, is one of the earliest markers of NC development, as it functions to specify the neural plate border and define the cells that will be competent to form the NC cell population (Basch et al., 2006; Khudyakov and Bronner-Fraser, 2009). SOX9, a high-mobility-group (HMG) domain-containing transcription factor, is a marker of prospective and early migratory NC cells (Basch et al., 2006). SNAI2 is a zinc-finger transcription factor that is highly conserved among vertebrate species and known to play an important role in EMT, functioning in a regulatory loop with Cadherin-6B during delamination (Schiffmacher et al., 2016; Taneyhill et al., 2007). SOX10, an SRY-related HMG-box family transcription factor, is necessary and sufficient to drive NC migration (Cheng et al., 2000; Honore et al., 2003; McKeown et al., 2005). Multiple studies have utilized analyses of gene expression via *in situ* hybridization and RNA-sequencing to characterize the timing of NC cell development in multiple species (Khudyakov and Bronner-Fraser, 2009; Soldatov et al., 2019; Williams et al., 2019). However, a comprehensive analysis of protein expression between species and across NC development is necessary to provide a framework for detailed functional analysis in the future.

Work from multiple organisms identified that NC GRN factors undergo post-transcriptional (Hutchins and Bronner, 2018) and post-translational (Hausser et al., 2019; Lander et al., 2011; Lee et al., 2012) modifications. Therefore, it is necessary to combine transcriptional analyses with protein expression studies to form a more complete picture of NC development. Using three closely related avian species, we characterize the spatiotemporal expression of PAX7, SOX9, SNAI2, and SOX10. Some key differences appear in quail, including earlier expression of the NC specifiers, SOX9, SNAI2, and SOX10, earlier NC cell migration out of the neural tube, and differences in cell migration modes. Further, NC EMT timing and mechanisms differ between the three avians. These results suggest that there are several key differences in the timing of NC cell development between species, emphasizing the need for functional *in vivo* protein studies in the future.

## 2. MATERIAL AND METHODS

### Avian Embryos

Fertilized chicken and quail eggs were obtained from UC Davis Hopkins Avian Facility and incubated at 37°C to the desired stages according to the criteria of Hamburger and Hamilton (HH)(Hamburger and Hamilton, 1951). Fertilized peafowl eggs were gifted from Dr. Pauline Perez. Fertilized eggs were incubated for different amounts of time to reach specific Hamburger Hamilton (HH)-equivalent stages, with quail being the fastest developing, followed by chick, then peafowl. To reach HH8, quail embryos were incubated 26- 29 hours, chick 28- 32 hours, and peafowl 36- 42 hours. Embryos were prepared for cryosectioning by equilibrating in 5% sucrose for 30 minutes to 1 hour at room temperature, then were transferred to 15% sucrose overnight in 4°C. Embryos were incubated in 10% gelatin overnight at 37°C, flash frozen in liquid nitrogen and stored at −80°C prior to sectioning. Embryos were then cyrosectioned in 14- 16 μm thick sections at −27°C on a Microm NX70 cryostat.

### Immunohistochemistry

Immunohistochemistry (IHC) was performed as previously described (Manohar et al., 2020; Rogers et al., 2013). Embryos were fixed on filter paper in 4% paraformaldehyde (PFA) in phosphate buffer for 15minutes at room temperature. After fixation, embryos were washed in 1X TBS (500 mM Tris-HCl pH 7.4, 1.5 M NaCl, and 10 mM CaCl_2_) containing 0.1% Triton X-100 (TBST+ Ca^2+^). Embryos were incubated in blocking buffer (TBST + Ca^2+^ with 10% donkey serum) for 1 hour at room temperature. Primary antibodies were diluted in blocking buffer and incubated with embryos for 24- 48 hours at 4°C (Table 1). After incubation with primary antibodies, embryos were washed with TBST + Ca^2+^ and incubated with Alexa Fluor secondary antibodies diluted (1:500) in blocking buffer for 12- 24 hours at 4°C. Embryos were then washed with TBST + Ca^2+^ and post-fixed in 4% PFA for 1 hour at room temperature or 12- 24 hours at 4°C. All embryos were imaged in whole mount and section using Zeiss Imager M2 with Apotome capability and Zen optical processing software.

**Table 1.** Antibodies used in study

| Antibody target | Dilution | Species | Isotype | Immunogen | Source |
| --- | --- | --- | --- | --- | --- |
| SOX10 | 1:500 | Mouse | Mouse IgG2a | SOX10 fusion protein Ag23191 | Protein tech (66786-1-Ig) |
| SOX9 | 1:500 | Human | Rabbit IgG | KLH- conjugated linear peptide corresponding to the C-terminal sequence of human SOX9 | Millipore (AB5535) |
| PAX7 | 1:5- 1:10 | Chicken | Mouse IgG1 | Amino acids 418-427 | DSHB (PAX7) |
| SNAI2 | 1:250 | Human | Rabbit IgG | Whole protein | Cell signaling (9585S) |

### Imaging

Fluorescence images were taken using Zeiss Imager M2 with Apotome.2 and Zen software (Karl Zeiss). Whole mount embryos were imaged using a EC Plan-Neofluar 10x/0.30 WD=5.2 M27. Embryo sections were imaged using a Plan-Apochromat 20x/0.8 WD=0.55 M27.

### Neural tube size analysis

Neural tube size was measured using Adobe Illustrator. Vectors were drawn from the dorsal to the ventral sides of the neural tube and lateral to lateral to measure width of the neural tube. The distance measured was then converted into microns using the in-image scale bar. Graphs were generated using GraphPad 9. An ordinary one-way ANOVA statistical test was used to compare data sets.

### Cell migration and cell count analyses

Cell migration was measured using Adobe Illustrator. A line was drawn from the dorsal midline of the neural tube to the end of the last migrated cell for SOX9, SNAI2, and SOX10+ cells at stages HH9 and HH11. The distance measured was then converted into microns using the in-image scale bar. Graphs were generated using GraphPad 9. An unpaired t-test was used for statistical analysis. Cells were counted in FIJI using the Cell Counter plug-in and plotted using GraphPad 9. The percentage of co-positive cells was calculated by dividing the number of cells expressing one protein over the number of the same cells expressing another protein. An ordinary one-way ANOVA statistical test was used to compare data sets.

### Protein localization analysis

Overlay of protein expression in whole mount and sections were performed in FIJI by subtracting background and superimposing all embryos from that stage to demonstrate that protein expression is consistent between embryos within the same stages as in (Simoes-Costa and Bronner, 2015).

### Fluorescence intensity analysis

The medial-lateral fluorescence intensity was calculated in FIJI by drawing a segmented line either from ventrolateral to dorsal or dorsal to lateral neural tube at a line width of 200 and a distance of 200 microns. A fluorescence intensity graph was generated through the plot profile plug-in, which was subtracted from background. The fluorescence intensity was then normalized by dividing each point by the highest fluorescence intensity value.

### Protein sequence and domain analysis

Chick and quail DNA and amino acid sequences were obtained from the National Center for Biotechnology Information (NCBI) and Kyoto Encyclopedia of Genes and Genomes (KEGG). Peafowl DNA sequence contigs were obtained from the NCBI *Pavo Cristatus* genome sequencing and assembly (Accession PRJNA413288) (Dhar et al., 2019). Both quail and chick sequences were used as queries to identify orthologous NC GRN genes. Reassembled DNA contigs were translated using Expasy Translate software (Ison et al., 2013). Full length amino acid sequences were then input into the Simple Modular Architecture Research Tool (SMART) Program (Letunic and Bork, 2018; Letunic et al., 2021) to create domain architecture maps that were exported as svg files and annotated in Adobe Illustrator. Multiple sequence alignments and comparisons were performed using Jalview (Procter et al., 2021; Waterhouse et al., 2009), Clustal Omega, and MView (Madeira et al., 2019).

## 3. RESULTS

To define the similarities and differences in the spatial and temporal expression and localization of NC-specific GRN proteins, we performed immunohistochemistry (IHC) on chick, quail, and peafowl embryos at multiple HH-equivalent developmental stages, using antibodies to mark NC progenitors (PAX7) and definitive NC cells (SOX9, SNAI2, and SOX10). Further, we quantified the percentage of cells expressing each protein and the differences in cranial NC cell migration between chick, quail, and peafowl. Here we present a detailed analysis of NC cell protein localization from neurulation (HH7) to late migration (HH15). The novelty in this study lies in our focus on spatiotemporal protein expression as a complement to previous RNA-sequencing (Williams et al., 2019) and *in situ* hybridization studies focusing on gene expression (Khudyakov and Bronner-Fraser, 2009).

### PAX7 timing and localization is similar between chick, quail, and peafowl

To define the similarities and differences between NC protein expression in chick, quail, and peafowl we performed IHC for PAX7, a NC progenitor marker, from neurulation (HH7, 1- 3 somite stage, SS) to migration (HH11, 13- 15 SS) stages. Further, we created composite expression overlays by aligning multiple embryos to identify if the spatial patterning is consistent between individuals. In confirmation with previous gene and protein expression studies in chick (Basch et al., 2006; Khudyakov and Bronner-Fraser, 2009), PAX7 protein is expressed in neural plate border cells at HH7 (Fig. 1A- 1C’, 1SS) in, chick, quail, and peafowl embryos, and continues to be expressed as the neural tube fuses at HH8 (Fig. 1D-1F’, 4- 5 SS). There exists a moderate amount of mediolateral variability in the location of the neural plate border in quail embryos (Fig. 1B’), but the spatial localization of PAX7 appears consistent between the three organisms at HH8 (Fig. 1D-1F’, 4- 5 SS). Although the onset of protein expression is similar between the three organisms, neural tube closure proceeds faster in quail and occurs at HH8 (Fig. 1E, 4 SS). By HH9, PAX7+ NC cells have begun to migrate laterally out of the now fused dorsal midbrain for all three organisms (Fig. 1G- 1I’, 7 SS).

**Figure 1.**
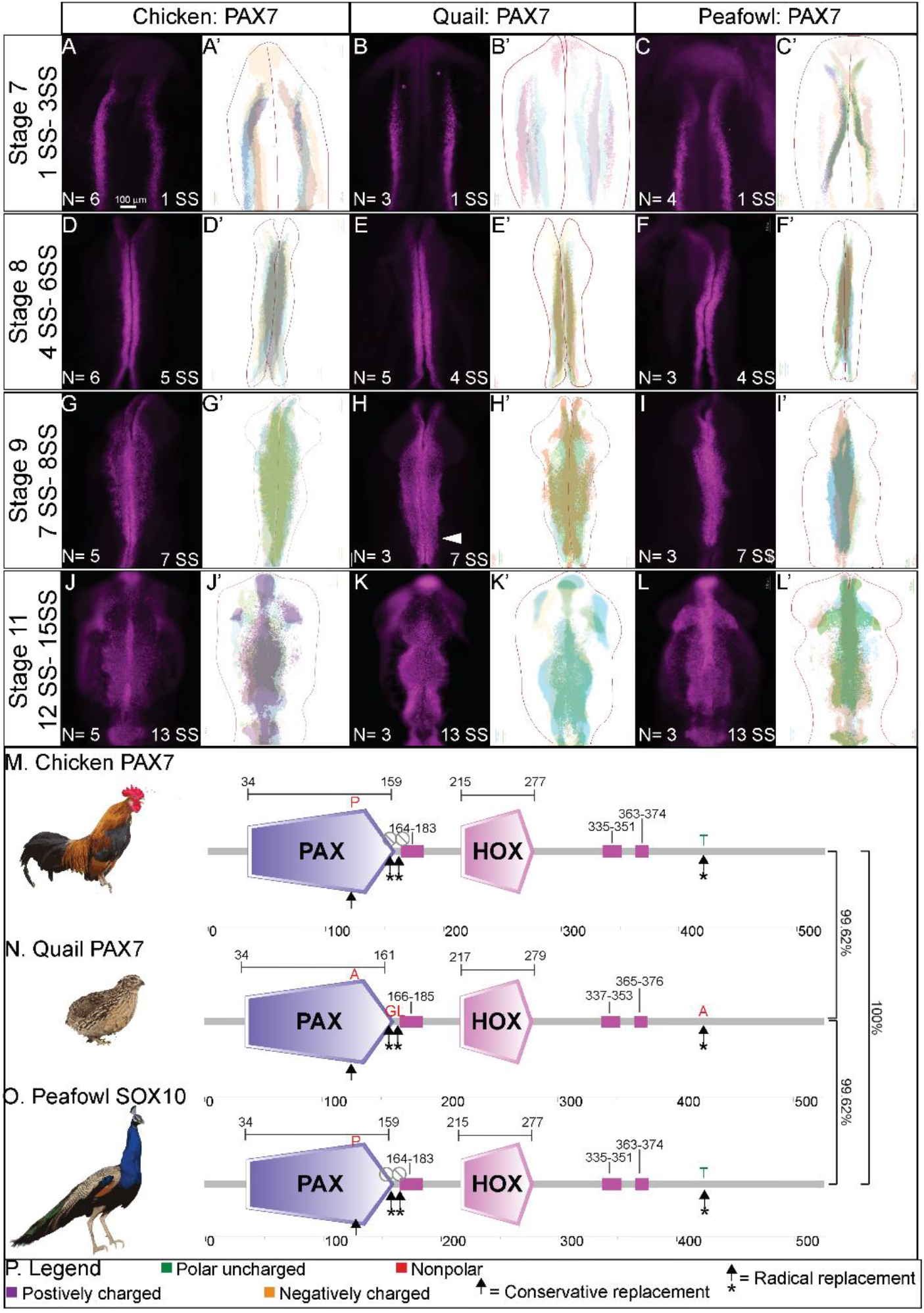
PAX7 expression timing in chick, quail, and peafowl whole embryos. IHC for PAX7 expression in (A-C’) HH7 (1 SS) neural plate border, (D-F’) HH8 (5 SS) dorsal neural tube, (G-I’) HH9 (7 SS) EMT stage NC cells, and (J-L’) HH11 (13 SS), neural tube and migratory NC cells. (A, D, G, J) in chick, (B, E, H, K) in quail, and (C, F, I, L) in peafowl embryos. (A’, B’, C’) schematic overlays of multiple embryos. Quail exhibits more posterior midline PAX7+ expression at HH9 (H, arrowhead). Number of embryos analyzed at each stage and used for schematic overlays indicated in IHC panels. Scale bar is 100 μm and all images were taken at the same magnification. (M-O) Amino acid sequences were aligned and compared, then analyzed in SMART to obtain domain diagrams. PAX (purple) and HOX (pink) domains are indicated on images and small pink boxes are low complexity domains. (M, O) Chick and peafowl PAX7 proteins are identical and (N) quail PAX7 has two amino acid replacements, one conservative (same type of amino acid, black arrow), one radical replacement (black arrow with asterisk), and two additional amino acids (black arrows with asterisks). (P) Legend for (M-O). Purple is positively charged amino acid, green is polar uncharged, yellow is negatively charged, red is nonpolar.

Some key differences appear at HH9 between species. In quail, PAX7+ NC cells appear to migrate in dense collectives compared to the chick, in which cells appear dense in the midline but sparser laterally (compare Fig. 1G to 1H). In peafowl, PAX7+ cells appear to have delayed migration out of the midline (compare Fig. 1I to Fig. 1G and 1H). Analysis of fluorescence intensity indicates similar PAX7 levels in the dorsal neural tube of chick and quail at HH8, and lower fluorescence intensity in peafowl (Supp. Fig. 1A- 1D, mean intensity 0.836, 0.787, and 0.298, data point 2 for chick, quail, and peafowl respectively), but higher levels in the quail and peafowl dorsal neural tube at HH9 (Supp. Fig. 1E- 1H, mean intensity 0.377, 0.673, and 0.539 for chick, quail, and peafowl, data point 2) and HH11 (Supp. Fig. 1I- 1L, mean intensity 0.352, 0.546, and 0.675 for chick, quail, and peafowl, data point 2) compared to chick (Supp. Fig. 1A- 1M). Further, the laterally migrating cells extend more posteriorly in quail (Fig. 1H, white arrow). At HH11, PAX7 expression remains strong in the midline of chick embryos, suggesting continued migration out of the neural tube (Fig. 1J, 1J’, 13 SS). However, migratory PAX7+ cells have migrated ventrally out of the dorsal focus, suggesting that the remaining PAX7+ cells are fated to remain in the dorsal neural tube (Fig. 1K, 1K’). In peafowl, PAX7+ cells are largely still close to the midline, suggesting delayed migration out of the neural tube (Fig. 1L and 1L’).

To begin to understand similarities and differences in PAX7 proteins between species, we performed multiple sequence alignment of the full-length chick, quail, and peafowl amino acid sequences. Chick and quail genomes are sequenced, annotated, and available on NCBI and KEGG, while the peafowl genome was reassembled using the available genome in NCBI (Dhar et al., 2019). As identified by Dhar et al., chick and Indian peafowl genomes are similar and the two are closely related (Dhar et al., 2019). Multiple sequence alignment identified that the PAX7 protein sequence is 100% conserved between chick and peafowl (Fig. 1M, 1O, Supp. Fig. 1M), while both sequences share 99.62% sequence identity with quail (Fig. 1N). The two sequences have 2 amino acid substitutions, one conservative (proline to alanine) and one radical (threonine to alanine) from chick and peafowl to quail (Fig. 1M- 1P, arrows) and quail PAX7 has two amino acids inserted in the N-terminal region glycine and leucine) that chick and peafowl are lacking (Fig. 1N).

To further compare PAX7 timing and localization, embryos were sectioned in the transverse plane and imaged at the midbrain axial level. At HH7 (3 SS), PAX7 marks the neural plate border cells that will be competent to form NC prior to neural tube closure (Fig. 2A- 2C). Quail and peafowl neurulation appears to occur at a slightly faster rate than in chick. As the neural tube fuses at HH8 (5 SS), chick and peafowl NC remain premigratory within the dorsal neural tube (Fig. 2D, 2 F) while quail cells appear to migrate laterally out of the closing neural tube (Fig. 2E). At HH9 (7 SS), PAX7+ premigratory NC progenitors remain in the dorsal neural tube in all organisms, and NC cells migrate dorsolaterally in chick (Fig. 2G), and dorsally in quail (Fig. 2H) and peafowl (Fig. 2I). Uniquely, peafowl NC cells are less densely packed at 7 SS compared to quail and chick. As migration continues at HH11 (13 SS), PAX7 expression is reduced in the leading cells at ventrolateral edge of the NC population in all organisms, but it remains expressed in the dorsal neural tube (Fig. 2J- 2L).

**Figure 2.**
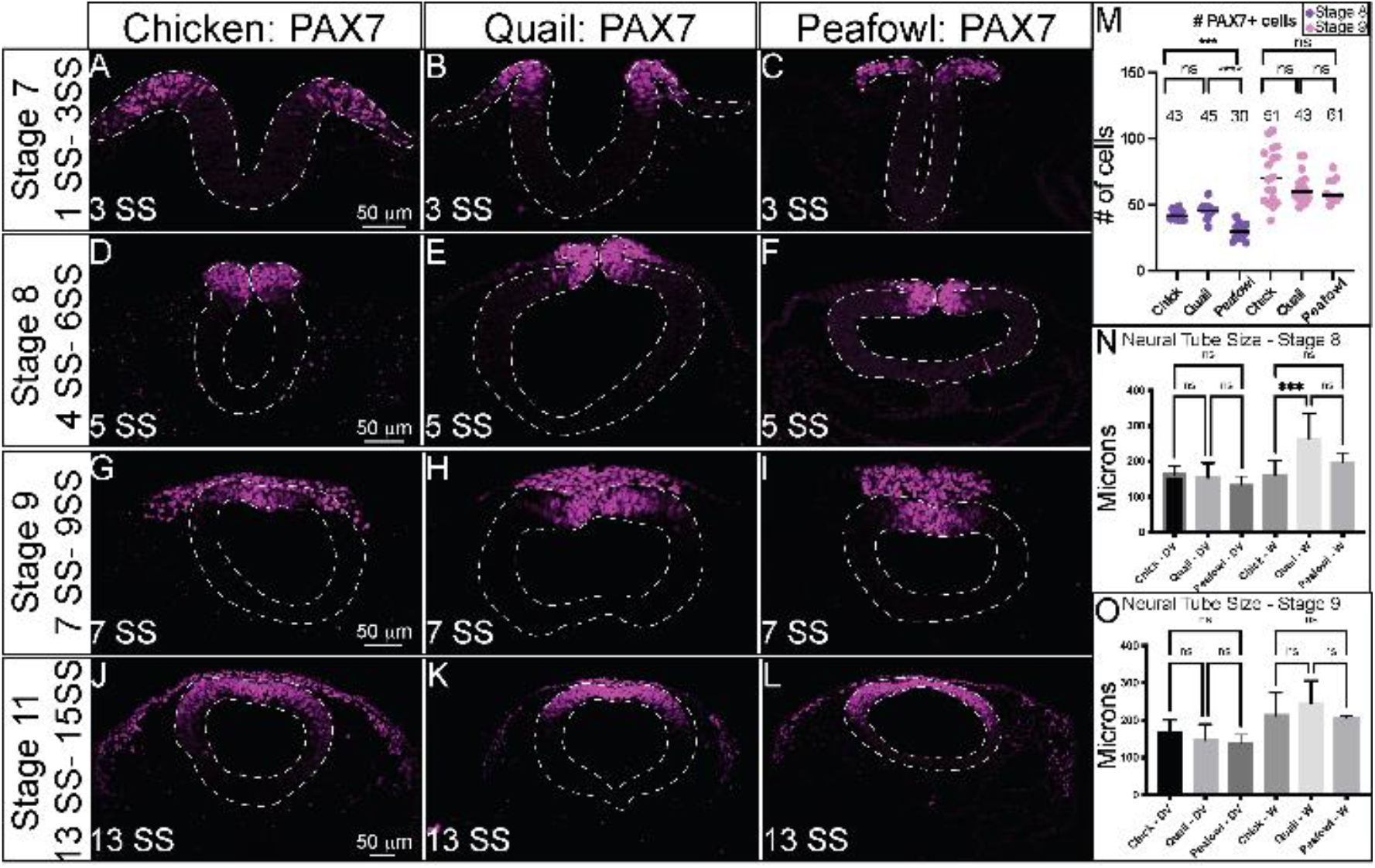
PAX7 expression timing in chick, quail, and peafowl sections. Transverse sections showing IHC for PAX7 in multiple stages of chick, quail, and peafowl embryos in (A-C) HH7, (D-F) HH8, (G-I) HH9, and (J-L) HH11. Dorsal is to the top and ventral is to the bottom. PAX7 expressed in all organisms by HH7 in the neural plate border (A- C), remains in the dorsal neural tube during neurulation (D-F), is expressed in premigratory and migratory NC cells (G- I), and is maintained in the dorsal neural tube and migratory NC cells (J-K). Scale bars are 50 μm and are as marked in first panel of each row. (M) The number of PAX7+ cells were quantified for HH8 and HH9, p = 0.001 between quail and peafowl, and p = 0.0002 between chick and peafowl, but no statistical difference was observed between chick and quail at HH8 (p = 0.990) or between any species at HH9 (p = 0.654 for chick and quail, p = 0.675 for chick and peafowl, and p = 0.999 for quail and peafowl). Ordinary one-way ANOVA statistical test used. (N) Neural tube size comparison at HH8. Chick dorsal-ventral (DV) compared to quail or peafowl DV is not significant, but width of the quail neural tube is larger than chick. Ordinary one-way ANOVA statistical test used (DV, p = 0.9756 and W, p = 0.002). (O) Neural tube size comparison at HH9. The DV and width of the neural tubes are no longer statistically significant. Ordinary one-way ANOVA statistical test used (DV, p = 0.3783 and W, p = 0.7398).

Given the differences in neural tube closure, we sought to determine if there were any differences in the number of PAX7+ cells in chick, quail, and peafowl at pre- and post-migratory stages (HH8 and HH9). However, at HH8, there is no statistical difference between the number of PAX7+ cells between chick and quail, however, there are significantly fewer cells in the peafowl compared to the chick (Fig. 2M, p= 0.0002) and quail (Fig. 2M, p= 0.001). By HH9, there is no statistical difference in the number of PAX7+ NC cells any of the organisms (Fig. 2M). In addition, we measured the differences in neural tube dorsal-ventral length and width between the three organisms, noting that the quail cranial neural tube appeared larger than chick. We determined that at HH8, the width of quail neural tube is significantly larger than in chick (Fig. 2N, p= 0.0002), but by HH9, the differences in width are no longer significant (Fig. 2K, p= 0.4). For both HH8 and HH9, we did not observe statistically significant differences in dorsal-ventral lengths. We noted that in quail embryos, the larger neural tubes at HH8 were followed by more collective NC cell migration at HH9, raising interesting questions regarding differences in NC migration mechanisms in quail. To better understand these cell migration variances, we analyzed the expression of EMT inducers, SOX9 and SNAI2, and migratory distances of cells positive for these proteins *in vivo* between the three species.

### SNAI2 is expressed early in the neural plate border in quail

*SNAI2* gene expression has been identified as early as HH8 and HH6.5 in chick and quail, respectively, with expression observed in the dorsal neural folds and neural plate border at the putative mid- and hindbrain levels (Basch et al., 2006; Khudyakov and Bronner-Fraser, 2009; Sakai et al., 2006). To determine if the protein localization in chick, quail, and peafowl was similar to its reported expression in chick at late HH7 (3 SS) (Taneyhill et al., 2007), we performed IHC for SNAI2 in various stages of these three avians. We observed SNAI2 protein in the dorsal neural folds as early as HH7 for all organisms, however, SNAI2 was not strongly expressed in chick and peafowl until 3-5 SS in the dorsal neural folds (Fig. 3A- 3A’, 3C, 3C’) while quail SNAI2 was expressed earlier, by 1 SS (Fig. 3B, 3B’, Fig. 4B). At HH8 (4- 6SS), SNAI2 is expressed in the fusing midbrain neural tube in chick, quail, and peafowl, but quail and peafowl SNAI2 expression extends more anteriorly into the diencephalon (Fig. 3B, 3C, 3D, 3E’, white arrows). At HH9 (7SS), SNAI2+ cells are undergoing EMT in all three organisms (Fig. 3G–I’). Whereas quail NC cells strongly express SNAI2 in the midline, chick and peafowl SNAI2+ cells appear more uniformly distributed (compare Fig. 3G, 3I to 3H). At HH11, SNAI2 is expressed in the midline premigratory NC cells and in migratory cells in all organisms, but peafowl development remains slower (Fig. 3J- 3L’, 13 SS). At HH11, SNAI2+ migratory cells have not extended as far ventrolaterally or anteriorly in peafowl compared to chick and quail (Fig. 3L, 3L’, asterisk). The SNAI2 amino acid sequences are highly conserved in all three organisms (Fig. 3M- 3P, Supp. Fig. 2M). Specifically, the amino acid sequences of chick and peafowl are 100% similar, and those sequences are 99.63% similar to the quail sequence, with a single conservative amino acid change on the N-terminus from glutamic acid to aspartic acid from chick and peafowl to quail (compare Fig. 3M, 3O to 3N).

**Figure 3.**
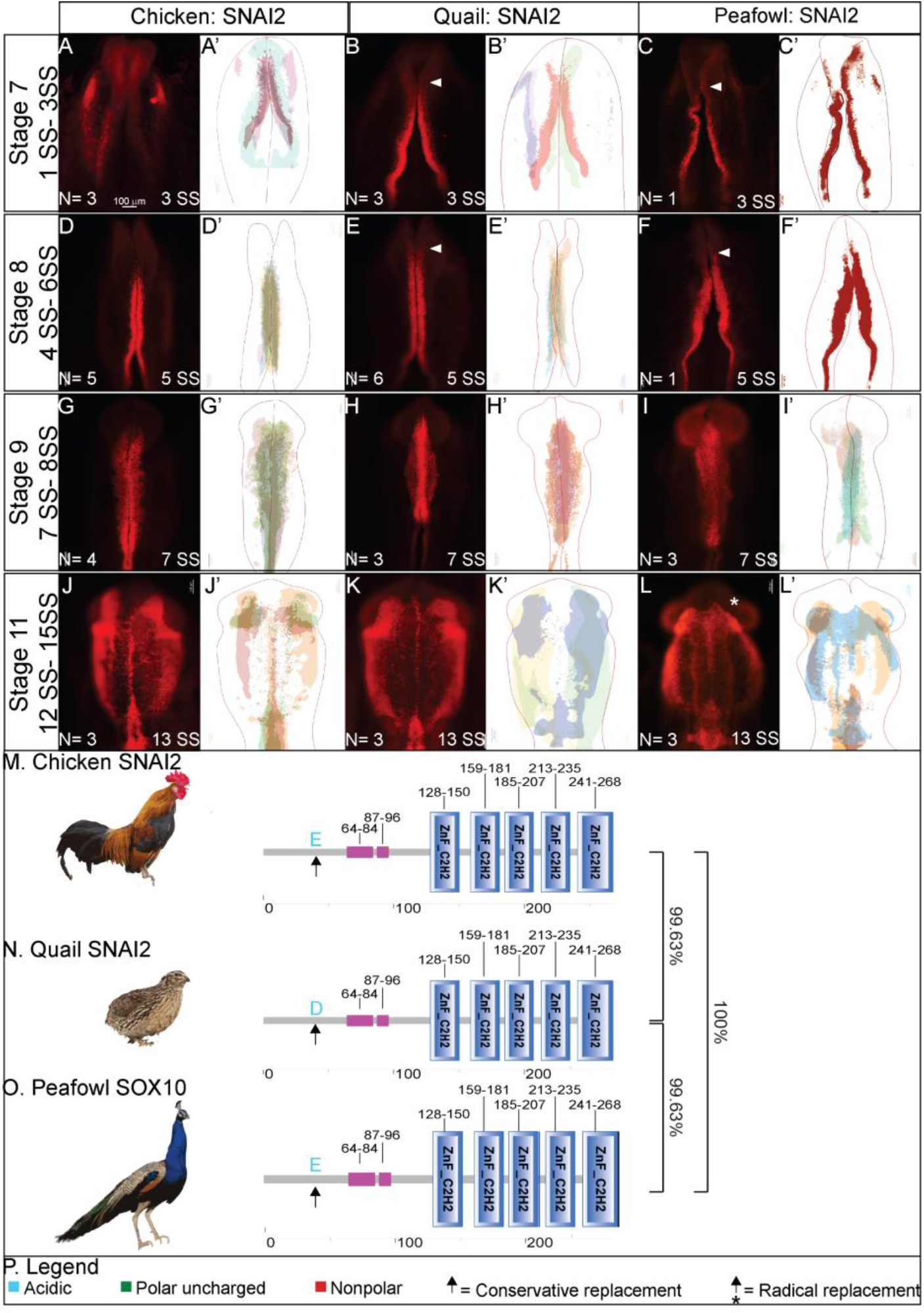
SNAI2 expression timing in chick, quail, and peafowl whole embryos. IHC for SNAI2 expression in (A-C’) HH7 (1 SS) neural plate border, (D-F’) HH8 (5 SS) dorsal neural tube, (G-I’) HH9 (7 SS) EMT stage NC cells, and (J-L’) HH11 (13 SS), neural tube and migratory NC cells. (A, D, G, J) in chick, (B, E, H, K) in quail, and (C, F, I, L) in peafowl embryos. (A’, B’, C’) schematic overlays of multiple embryos. Quail and peafowl exhibit more anterior SNAI2+ expression than chick between HH7 to HH8 (compare A, D to B, E and C, F, arrowhead). Number of embryos analyzed at each stage and used for schematic overlays indicated in IHC panels. Scale bar is 100 μm and all images were taken at the same magnification. (M- O) Amino acid sequences were aligned and compared, then analyzed in SMART to obtain domain diagrams. Zinc finger domains (blue) and small pink boxes are low complexity domains are shown on schematics. (M, O) Chick and peafowl SNAI2 proteins are identical and (N) quail SNAI2 has one conservative amino acid replacement (same type of amino acid, black arrow). (P) Legend for (M-O). Blue is acidic amino acid, green is polar uncharged, red is nonpolar.

**Figure 4.**
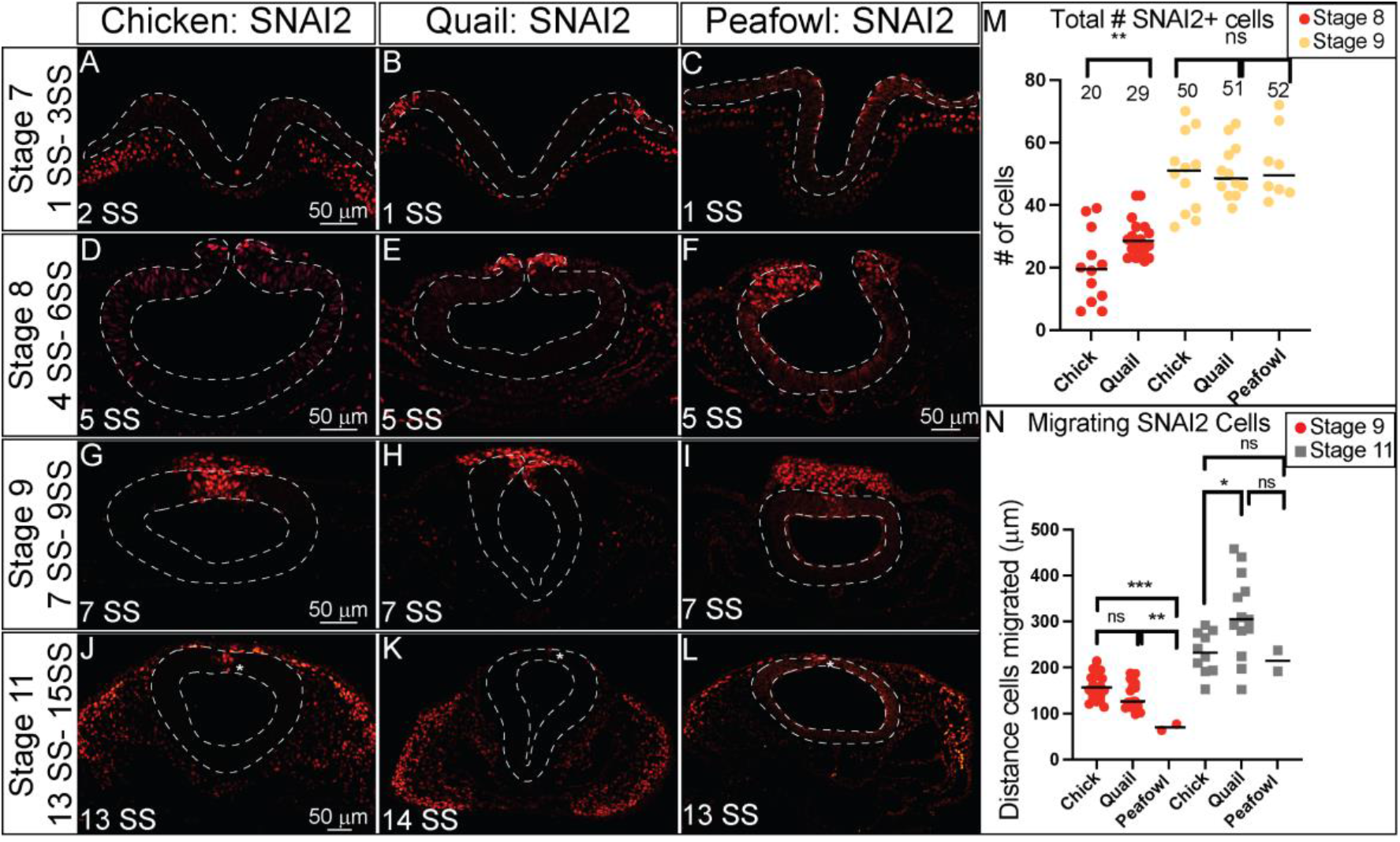
SNAI2 expression timing in chick, quail, and peafowl sections. Transverse sections showing IHC for SNAI2 in multiple stages of chick, quail, and peafowl embryos in (A-C) HH7, (D-F) HH8, (G-I) HH9, and (J-L) HH11. Dorsal is to the top and ventral is to the bottom. In quail, SNAI2 is expressed at HH7 in the neural plate border (compare B to A and C), is expressed in the dorsal neural tube during neurulation/NC specification at HH8 in all three organisms (D-F), is expressed in premigratory and migratory NC cells at HH9 during EMT (G- I), and is maintained in the dorsal neural tube in a small number of premigratory NC cells but is highly expressed in migratory NC cells at HH11 (J- K). Scale bars are 50 μm and are as marked in first panel of each row. (M) The number of SNAI2+ cells were quantified for HH8 and HH9, at HH8 quail had statistically more cells than chicken, but by HH9 that statistical significance was no longer observed. Ordinary one-way ANOVA statistical test was used (HH8, p = 0.0571 and HH9, p = 0.9975). (N) Cell migration was measured from the neural tube midline to the furthest migrated cell for HH9 and HH11. At HH9, chicken and quail cells migrated further than peafowl (p = 0.004 chick and peafowl and p =0.008 for quail and peafowl, p = 0.052 for chick and quail). At HH11, quail cells have migrated further than chick (p = 0.043 for chick and quail, p = 0.946 for chick and peafowl, and p = 0.209 for quail and peafowl).

In transverse section, it is clear that SNAI2 expression in quail (Fig. 4B, HH7, 1 SS) precedes that of chick (Fig. 4A, HH7, 2 SS) and peafowl (Fig. 4C, HH7, 1SS) in the neural plate border. At HH8 (5SS), quail SNAI2+ cells are starting to undergo EMT or migrate out of the neural tube while chick and peafowl cells remain within the neuroepithelium (Fig. 4D-4F, 5 SS). As with previous markers, at HH9 (7 SS), quail SNAI2+ cells appear to migrate more collectively during EMT compared to chick, while the chick embryos maintain stronger SNAI2 expression in the premigratory population than either quail or peafowl (Fig. 4G- 4I). By HH11 (13-14 SS) SNAI2 is expressed in migrating NC cells in chick, quail, and peafowl with very few SNAI2+ premigratory NC cells (Fig. 4J- 4L, asterisk). As SNAI2 drives NC EMT (Taneyhill et al., 2007) and is expressed earlier in quail, we analyzed the number of total SNAI2+ cells and the distance that SNAI2+ cells migrated from the midline. There were significantly more quail SNAI2+ NC cells at HH8 in quail embryos compared to chick, but by HH9, this difference was lost (Fig. 4M). We compared the distance that SNAI2+ cells migrated from the midline to the leading edge in all three organisms, and at HH9, both chick and quail cells migrated farther ventrolaterally than peafowl, but by HH11, the distance between chick or quail with peafowl was no longer significantly different (Fig. 4N, p = 0.004 for chick and peafowl and p =0.008 for quail and peafowl, p = 0.052 for chick and quail at HH9). However, at HH11 quail SNAI2+ NC cells migrated further than chick SNAI2+ cells (Fig. 4N, p = 0.043).

Fluorescence intensity analyses demonstrated higher intensity for SNAI2+ cells in quail premigratory NC cells as the mean intensity for chick was 0.449 compared to 0.827 for quail at the leading edge (Supp. Fig. 2A- 2C, data point 2). Early migratory NC cells had a mean intensity of 0.372 in chick, 0.611 in quail, and 0.774 in peafowl (Supp. Fig. 2D-2G, data point 2), but similar intensity between species for migrating NC cells (mean intensity 0.451 for chick, 0.573 for quail, and 0.679 for peafowl) (Supp. Fig. 2H- 2K, data point 2).

### SOX9 is more robustly expressed in quail and peafowl during specification

The *Sox9* gene is expressed at HH8 in chick in the closing neural folds (Basch et al., 2006; Betancur et al., 2009; Yardley and Garcia-Castro, 2012) and as early as HH6 in the anterior neural folds of quail embryos (Sakai et al., 2006). To identify whether the protein expression mirrors the spatiotemporal gene expression, we performed IHC for SOX9 in multiple stages of chick, quail, and peafowl embryos. In contrast to PAX7, SOX9 protein was not expressed prior to late HH7 in any of the three species. Chick SOX9 is barely apparent in the dorsal neural tube at late HH7 in whole mount (Fig. 5A, 5A’, 3 SS). In contrast, at HH7, quail and peafowl SOX9 expression is visible in several NC cells within the closing neural tube (Fig. 5B- 6C’, 3 SS). At HH8, SOX9 is expressed in the dorsal neural tube at the midbrain level and is more robust in the quail and peafowl (Fig. 5D-5F’, 5 SS). At HH9 during EMT, SOX9+ cells are expressed in migrating NC cells (Fig. 5G- I’, 7 SS). Similar to PAX7 expression, chick SOX9+ cells appear more dispersed during migration than quail or peafowl SOX9+ cells, which appear more condensed in quail and retained to the midline in peafowl (compare Fig. 5G to 5H and 5I). Additionally, peafowl SOX9+ cells have not migrated as far as those of chick and quail at this stage. By HH11, SOX9 is still expressed in migrating NC cells, as well as the otic placode in all species (Fig. 5J- 5L’, 13 SS). Few SOX9+ cells remain in the neural tube at this stage (Fig. 5J, 5K, 5L white arrows), but peafowl SOX9 cells are more dorsally located than either chick or quail, and similar to SNAI2, SOX9+ cells do not migrate as far anteriorly in peafowl at this stage (Fig. 5L, 5L’). As with the PAX7 and SNAI2 sequence alignments, chick and peafowl SOX9 amino acid sequences are 100% conserved and are 99.39% similar to the quail SOX9 sequence (Fig. 5M- 5O, Supp. Fig. 3M). There are both conservative (serine to asparagine) and radical replacements (lysine to glutamic acid and alanine to threonine) in the sequences as well as a lost amino acid (glutamine) in chick and peafowl compared to quail (Fig. 5M- 5P).

**Figure 5.**
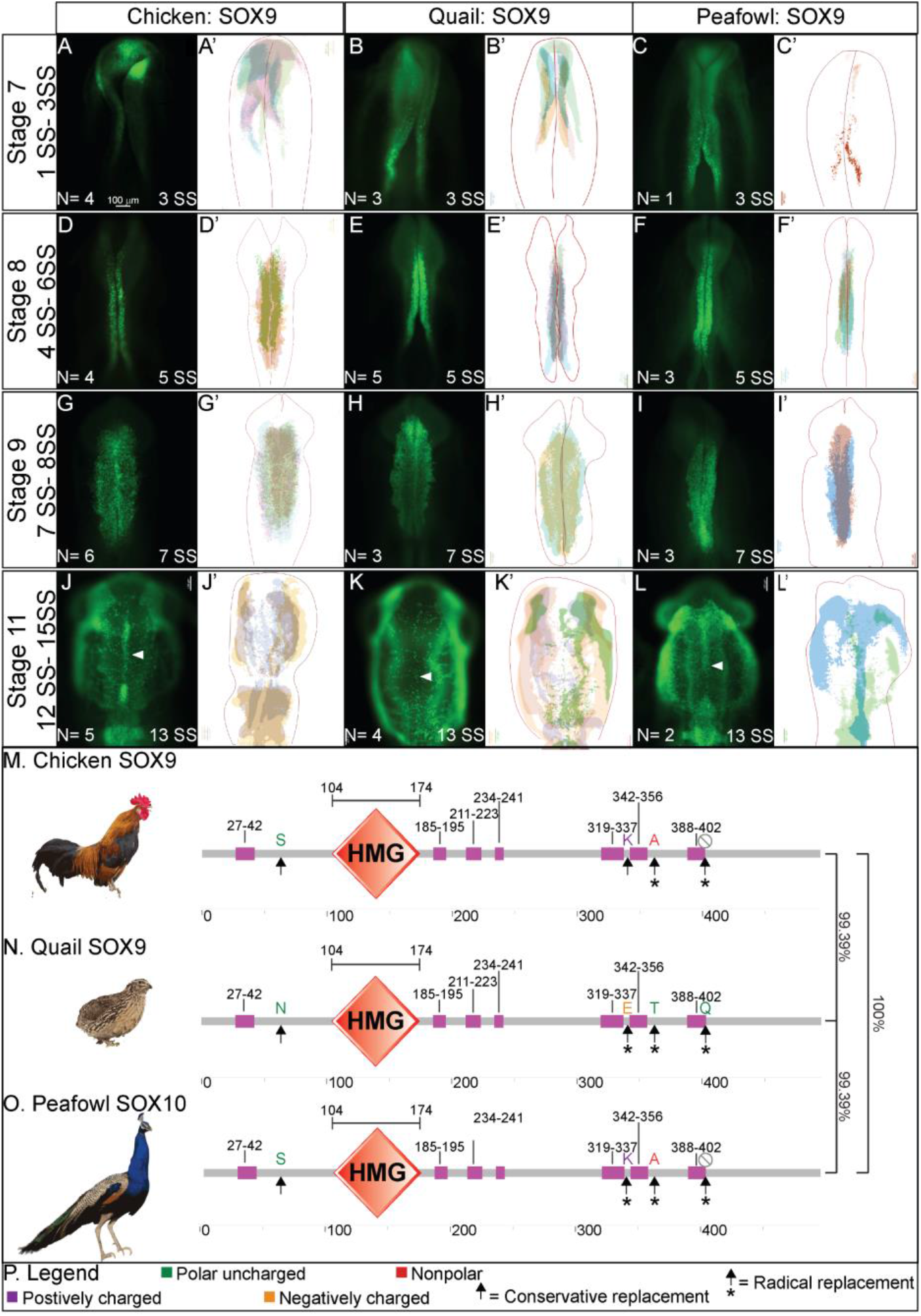
SOX9 expression timing in chick, quail, and peafowl whole embryos. IHC for SOX9 expression in (A-C’) HH7 (1 SS) neural plate border, (D-F’) HH8 (5 SS) dorsal neural tube, (G-I’) HH9 (7 SS) EMT stage NC cells, and (J-L’) HH11 (13 SS), neural tube and migratory NC cells. (A, D, G, J) in chick, (B, E, H, K) in quail, and (C, F, I, L) in peafowl embryos. (A’, B’, C’) schematic overlays of multiple embryos. Chick and peafowl maintain more SOX9+ cells in premigratory cells at HH11 than quail (compare J and L to K, arrowhead). Number of embryos analyzed at each stage and used for schematic overlays indicated in IHC panels. Scale bar is 100 μm and all images were taken at the same magnification. (M- O) Amino acid sequences were aligned and compared, then analyzed in SMART to obtain domain diagrams. HMG domains (orange) and low complexity domains (pink) are shown in diagram. (M, O) Chick and peafowl SOX9 proteins are identical and (N) quail SOX9 has four amino acid replacements, one conservative (same type of amino acid, black arrow), two radical replacements (black arrow with asterisk), and one additional amino acid (black arrows with asterisks). (P) Legend for (M-O). Purple is positively charged amino acid, green is polar uncharged, yellow is negatively charged, red is nonpolar.

**Figure 6.**
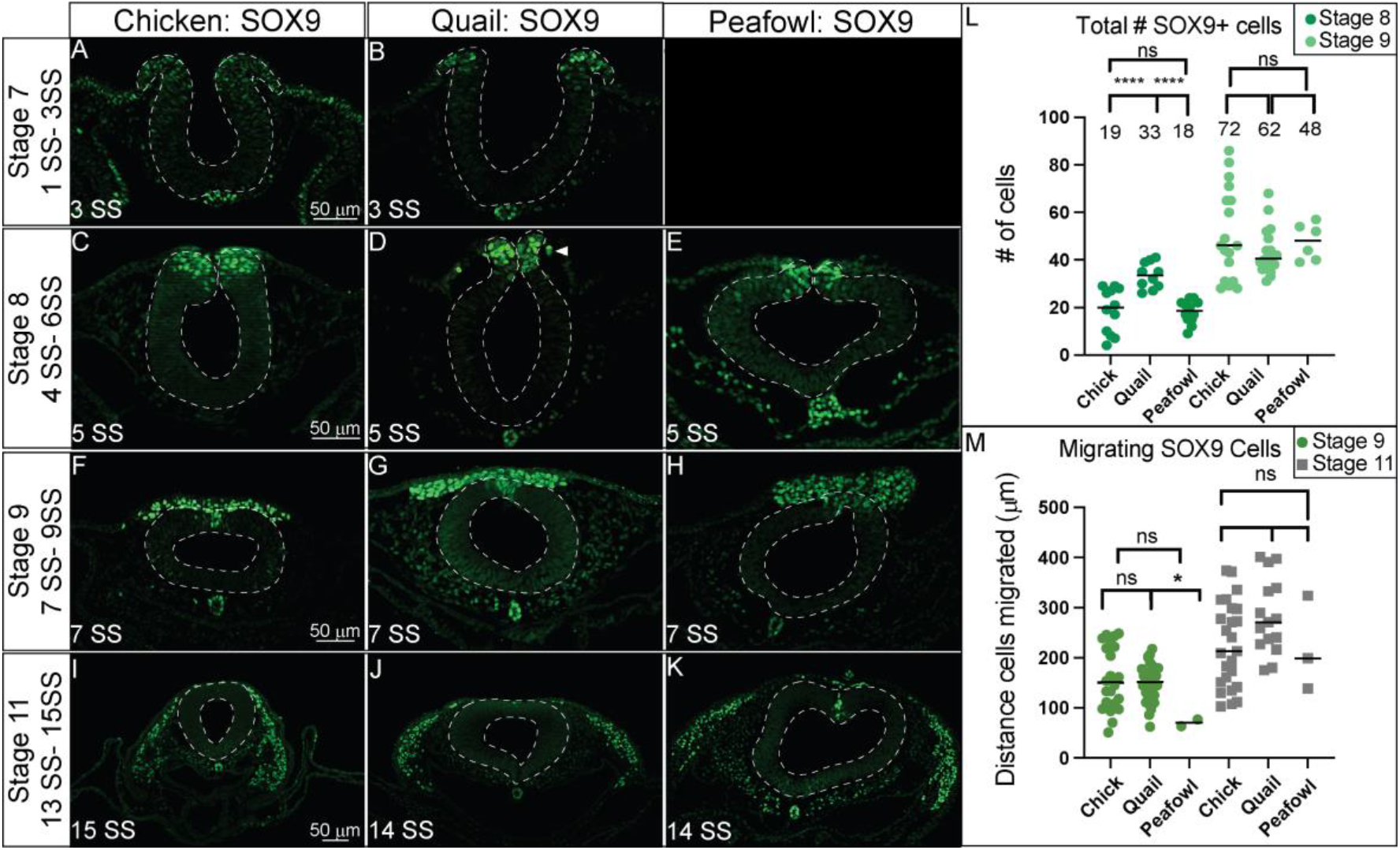
SOX9 expression timing in chick, quail, and peafowl sections. Transverse sections showing IHC for SOX9 in multiple stages of chick, quail, and peafowl embryos in (A, B) HH7 (not obtained for peafowl), (C- E) HH8, (F- H) HH9, and (I- K) HH11. Dorsal is to the top and ventral is to the bottom. SOX9 is expressed in quail at HH7 in the neural plate border/dorsal neural tube (compare B to A), but we were unable to obtain peafowl sections at this stage to compare. SOX9 is expressed in the dorsal neural tube during neurulation/NC specification at HH8 in all three organisms (C- E), is expressed in premigratory and migratory NC cells at HH9 during EMT (F- H), and is maintained in the dorsal neural tube in a small number of premigratory NC cells (most prevalent in peafowl), but is highly expressed in migratory NC cells at HH11 (I- K). Scale bars are 50 μm and are as marked in first panel of each row. (L) The number of SOX9+ cells were quantified for HH8 and HH9. At HH8, quail expressed more SOX9+ cells than chick (p = 0.0001) and peafowl (p = 0.0001), but by HH9 this difference was no longer statistically significant (p = 0.997). Ordinary one-way ANOVA statistical test used. (M) Cell migration was measured from the neural tube midline to the furthest migrated cell for HH9 and HH11. No statistical significance was observed, except between quail and peafowl at HH9 (p = 0.0373).

In sections, chick SOX9 is expressed at very low levels in the dorsal neural folds while quail SOX9 is upregulated in premigratory NC cells by late HH7 (Fig. 6A, 6B, 3 SS). Due to limited availability, we were unable to obtain a section in peafowl at this stage. SOX9 is also expressed in the notochord at all stages tested in all three species (Fig 6A- 6K). Similar to PAX7+ cells, quail SOX9+ NC cells appear to begin migrating at the onset of neural tube closure at HH8 (5 SS, arrowhead) while chick and peafowl cells remain neuroepithelial (Fig. 6C- 6E). We confirmed that quail cells migrate more collectively at HH9 as they are more densely packed compared to chick and peafowl (Fig. 6F- 6H, 7 SS). At HH11, most chick and quail SOX9+ cells have migrated out of the neural tube while peafowl still has some premigratory cells remaining (Fig. 6I- 6K, 14 -15 SS).

To determine if the timing of NC specification by SOX9 expression was equivalent between species, we counted the total number of NC cells positive for SOX9 at stages HH8 and HH9 (Fig. 6L). We identified that there are an average of 19 SOX9+ NC cells in chick, 33 SOX9+ cells in quail, and 18 SOX9+ cells in peafowl at HH8 (4- 6 SS) (Fig. 6L, p= 0.0001 for chick and quail, p = 0.0001 for quail and peafowl, and p = 1.0 for chick and peafowl). At HH9, the average number of SOX9+ NC cells is no longer statistically different between organisms although there is variability in the total number of cells expressing SOX9 (Fig. 6L, p= 0.878 for chick and quail, p = 1 for quail and peafowl, and p = 1.0 for chick and peafowl). To identify if the differences in timing of quail NC specification lead to increased migratory distance, we measured the distance between the leading SOX9+ NC cells from the midline of the embryos. While there are no statistical differences in migration distance between chick and quail or chick and peafowl at HH9, quail SOX9+ cells migrated further compared to peafowl (Fig. 6J, p = 0.037). By HH11 there was no statistical difference in migration distance from the midline in any of the species suggesting that the early onset of specification does not necessarily lead to faster NC migration (Fig. 6J, p = 0.0982 for chick and quail, p = 0.9943 for chick and peafowl, and p = 0.463 for quail and peafowl). These results suggest that there may be differences in NC timing between avian species. Additional analysis of SOX9 fluorescence intensity in the dorsal neural tube at HH8 confirms that SOX9 levels in the dorsal neural tube of chick, quail, and peafowl are similar at HH8 (mean intensity for chick = 0.521, 0.875 for quail, and 0.617 for peafowl, data point 2) (Supp. Fig. 3A- 3D). However, at HH11, chick and peafowl SOX9 levels in the dorsal neural tube exhibited stronger intensity than quail (mean intensity for chick = 0.321, 0.167 for quail, and 0.264 for peafowl, data point 1l), while quail cells had stronger intensity in the migratory regions (mean intensity for quail = 0.400 and 0.246 for chick, data point 2) (Supp. Fig. 3I- 3L).

### SOX10 expression occurs just prior to EMT in all species

Previous analysis of *Sox10* gene expression in chick identified expression in the cranial neural folds at HH8+, in migrating NC cells at HH9, and in otic placodes at HH10+ (Basch et al., 2006; Betancur et al., 2010; McKeown et al., 2005). Quail *SOX10* gene expression has been reported as barely detectable at HH8 but is strongly expressed in the cranial neural folds and migrating NC cells by HH9 (Sakai et al., 2006). Here we compare SOX10 protein expression in chick, quail, and peafowl embryos. Neither organism has discernible SOX10 expression at HH7 (data not shown). Similar to *Sox10* gene expression, chick SOX10 protein is barely detectable at HH8, but quail and peafowl SOX10 is expressed more strongly in the dorsal neural folds (Fig. 7A- 7C’, 5 SS), which contrasts with previous reports that *Sox10* gene expression was not detected until HH9 in quail (Sakai et al., 2006). By HH9, SOX10 is expressed in early migrating NC cells in all three organisms (Fig. 7D-7F’). Chick SOX10+ cells are more dispersed while quail and peafowl SOX10+ cells appear more densely packed (Fig. 7D-7F’, 7 SS). By HH11, migrating NC cells continue to express SOX10 sparsely in the midline and more intensely in the ventrolateral NC populations (Fig. 7G- 7I’, 13 SS). SOX10 is also expressed in the otic placodes in all organisms at HH11 (Fig. 7G- 7I’). SOX10 protein expression is similar to its previously reported gene expression in chick with some differences in timing in quail. Of the proteins compared, the SOX10 amino acid sequence exhibited the most differences between species (Supp. Fig. 4M). In the case of SOX10, chick and peafowl are 98.92 similar (Fig. 7J, 7L), chick and quail are 80.40% similar (Fig. 7J, 7K), and quail and peafowl have 81.60% sequence similarity (Fig. 7K, 7L). The quail SOX10 amino acid sequence has 104 amino acids on the terminal end that do not appear to exist in either chick or peafowl sequences (Fig. 7J- 7L, Supp. Fig. 4M). There are two conservative amino acid replacements in the N-terminus (aspartic acid to glutamic acid and asparagine to serine) in chick and peafowl compared to quail (compare Fig. 7J, 7L to 7K). The remaining amino acid changes are derived from radical replacements. In the case of SOX10, the peafowl sequence mirrors the chick sequence and differs from quail at positions 108 (aspartic acid to glutamic acid), 137 (asparagine to serine), and 417 (threonine to alanine). The peafowl sequence mirrors the quail sequence and differs from chick at positions 139 (proline to serine), 143 (glycine to serine), 299 (alanine to serine), 307 (threonine to alanine). At position 142, chick and quail mirror each other and differ from peafowl (alanine to serine) (Fig. 7J- 7M, Supp. Fig. 4M). Although the protein timing differs and the localization is similar between organisms, future work will focus on defining if the protein functions are conserved between species.

**Figure 7.**
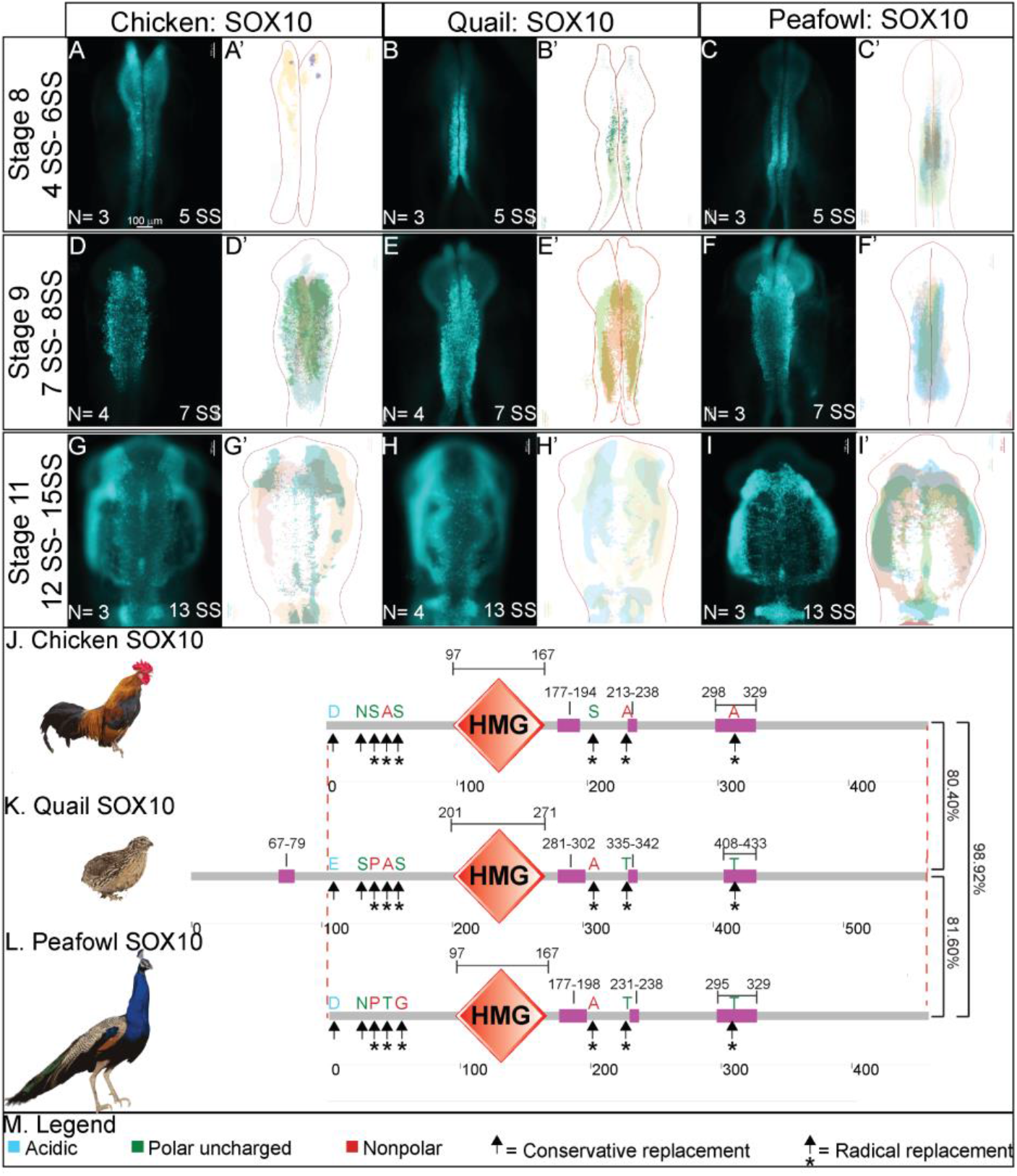
SOX10 expression timing in chick, quail, and peafowl whole embryos. IHC for SOX10 expression in (A-C’) HH8 (5 SS) dorsal neural tube, (D-F’) HH9 (7 SS) EMT stage NC cells, and (G-I’) HH11 (13 SS), neural tube and migratory NC cells. (A, D, G) in chick, (B, E, H) in quail, and (C, F, I) in peafowl embryos. (A’- C’, D’- F’, G’- I’) schematic overlays of multiple embryos. Peafowl embryos have delayed migration of SOX10+ cells compared to chick and quail as the cells remain visible in the dorsal focal plane (compare I to G and H). Number of embryos analyzed at each stage and used for schematic overlays indicated in IHC panels. Scale bar is 100 μm and all images were taken at the same magnification. (J- L) Amino acid sequences were aligned and compared, then analyzed in SMART to obtain domain diagrams. HMG domains (orange) and low complexity domains (pink) are shown on schematics. (J- L) Peafowl SOX10 differs from the other proteins profiled here. It shares amino acid sequences with both chick and quail (same type of amino acid, black arrow, different type of amino acid black arrow plus asterisk), and has one amino acid that is unique from both chick and quail, which resemble each other. (M) Legend for (J- L). Blue is acidic amino acid, green is polar uncharged, red is nonpolar.

In transverse section, SOX10 protein is expressed in a limited number of premigratory NC cells, colocalizing with a small proportion of the PAX7, SNAI2, and SOX9+ populations in all species after HH8 (Fig. 8). At late HH8, SOX10 appears localized to premigratory NC cells in the neural tube, but is expressed as early as 4 SS in quail (Fig. 8A- 8C, 5 SS). At HH9, very few cells in the neural tube are SOX10+ in chick, quail, or peafowl, while most migrating cells express SOX10 (Fig. 8D-8F, 7 SS). As with previous markers, SOX10+ migratory NC cells are more densely packed in quail compared to chick (Fig. 8D-8E). By HH11, SOX10 is expressed in migratory cells, and SOX10+ quail NC cells continue to migrate collectively into the ventrolateral regions of the head in contrast to the more mesenchymal chick and peafowl cells (Fig. 8G- 8I, 13 SS).

**Figure 8.**
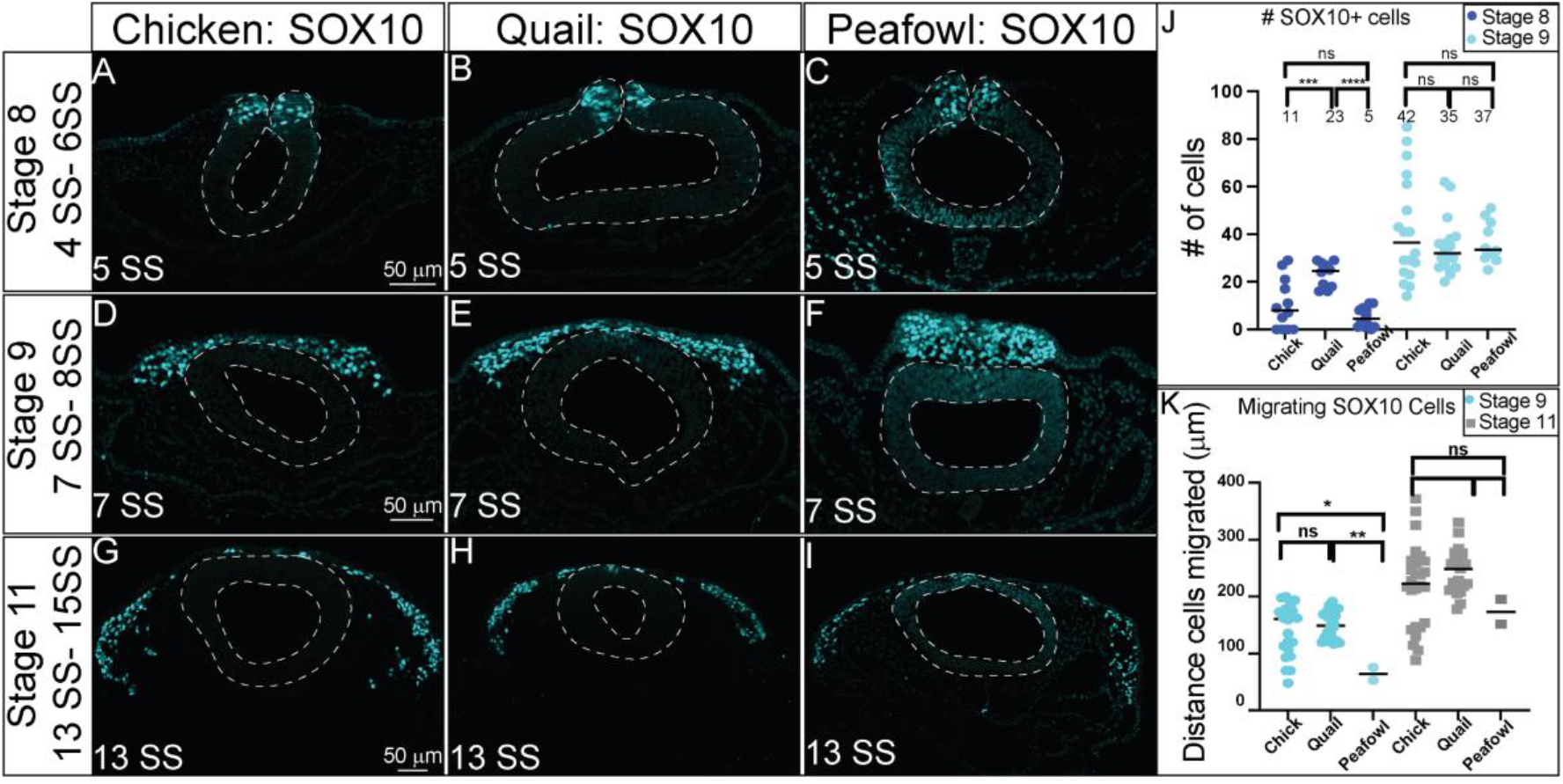
SOX10 expression timing in chick, quail, and peafowl sections. Transverse sections showing IHC for SOX10 in multiple stages of chick, quail, and peafowl embryos in (A-C) HH8, (D-F) HH9, and (G-I) HH11. Dorsal is to the top and ventral is to the bottom. SOX10 is expressed in the dorsal neural tube during neurulation/NC specification at HH8 in all three organisms (A- C), is expressed in premigratory and migratory NC cells at HH9 during EMT (D-F), and is maintained in the dorsal neural tube in a small number of premigratory NC cells but is highly expressed in migratory NC cells at HH11 (G- I). Scale bars are 50 μm and are as marked in first panel of each row. (J) The number of SOX10+ cells were quantified for HH8 and HH9. At HH8, quail expressed more SOX10+ cells than chick (p = 0.001) and peafowl (p = 0.001), but by HH9 this difference was no longer statistically significant (p = 0.990). An ordinary one-way ANOVA statistical test was used. (K) Cell migration was measured from the neural tube midline to the furthest migrated cell for HH9 and HH11. At HH9, chick (p = 0.019) and quail (p = 0.009) cells migrated further than peafowl, but no distance was observed between quail and chick (p = 0.760). At HH11 no statistical difference was observed between species (p = 0.288 for chick and quail, p = 0.556 for chick and peafowl, and p = 0.244 for quail and peafowl). An Ordinary one-way ANOVA statistical test was used.

To confirm developmental differences in NC timing at premigratory stages, we counted the number of SOX10+ cells at HH8 and 9. At HH8 (3- 5 SS), there were 2-fold more SOX10+ cells in quail embryos compared to chick and 4-fold more SOX10+ cells in quail embryos compared to peafowl (Fig. 8J, p= 0.0010 for chick and quail, and p = 0.0001 for quail and peafowl), but this difference was lost by HH9. At HH9, peafowl cells migrated a significantly shorter distance compared to either chick (p=0.0019) or quail (p= 0.009). In contrast to SNAI2+ cells, there were no differences in the distances migrated by SOX10+ NC cells in any species at HH 11 (Fig. 8K).

Fluorescence intensity measurements of premigratory SOX10 positive cells at HH8 in sectioned embryos showed higher intensity for quail and peafowl compared to chick, consistent with whole mount and section data (mean intensity for chick = 0.308, 0.665 for quail, and 0.8 for peafowl, data point 2) (Supp. Fig. 4A- 4D). However, by HH9, fluorescence intensity is similar between chick and quail (mean intensity for chick = 0.415 and 0.491 for quail, data point 2) and by HH11, the majority of SOX10+ cells have migrated out of the neural tube and are migrating laterally (mean intensity for chick = 0.429, 0.671 for quail, and 0.444 for peafowl, data point 2) (Supp Fig. 4E–4L).

### Heterogeneous protein expression identifies subsets of NC populations in all organisms

Prior work established that the premigratory NC population is multipotent (Bhattacharya et al., 2018; Buitrago-Delgado et al., 2015; Kerosuo et al., 2015) and that at the transcript level, multiple NC-related genes coding for transcription factors are expressed throughout the dorsal neural tube (Lignell et al., 2017; Williams et al., 2019). However, information defining the timing and localization of the NC proteins during NC specification and EMT is still lacking. To identify the extent of overlapping and discrete expression of the NC progenitor protein PAX7 with specifier/EMT driving proteins (SNAI2, SOX9, SOX10), we performed multiplexed IHC during specification (HH8, 4-6 SS) and EMT (HH9, 7-9 SS) stages. The number of NC cells expressing each protein was quantified in transverse sections from multiple embryos at equivalent HH stages (Figs. 9, 10). Comparing the total number of PAX7, SOX9, and SOX10+ cells in chick, quail, and peafowl at HH8 identified both similarities and differences in NC developmental timing between species. There are no significant differences in the number of PAX7+ cells in chick and quail at HH8, but there are significantly fewer PAX7+ cells in peafowl at HH8 compared to both chick and quail (Fig. 9A, 9C, 9G, 9K, chick to peafowl, p= 0.0002, quail to peafowl, p= 0.001). Further, as reported in figures 6 and 8, there were more SOX9 and SOX10+ NC cells in quail than either chick or peafowl at HH8 (Fig. 9D, 9E, 9H, 9I, 9L, 9M, p= 0.001 for quail compared to chick or peafowl for both markers). During specification in chick (HH8, 4- 6 SS), 48% and 24% of PAX7+ cells are also positive for SOX9 or SOX10, respectively (Fig. 9B, 9C- 9F). In quail, 77% and 51% of PAX7+ cells co-express SOX9 and SOX10, respectively at HH8 (4- 6 SS) (Fig. 9B, 9G- 9J). In peafowl embryos, 61% and 17% of PAX7+ cells co-express SOX9 and SOX10, respectively. These data suggest that there is a more rapid onset of NC specifiers in quail than chick or peafowl embryos with the NC program beginning at earlier HH stages. In contrast to the low to moderate co-expression of SOXE proteins with PAX7, SNAI2 appeared to be expressed in more NC cells at HH8 than the other specifiers in all three organisms (Fig. 9O- 9Z).

**Figure 9.**
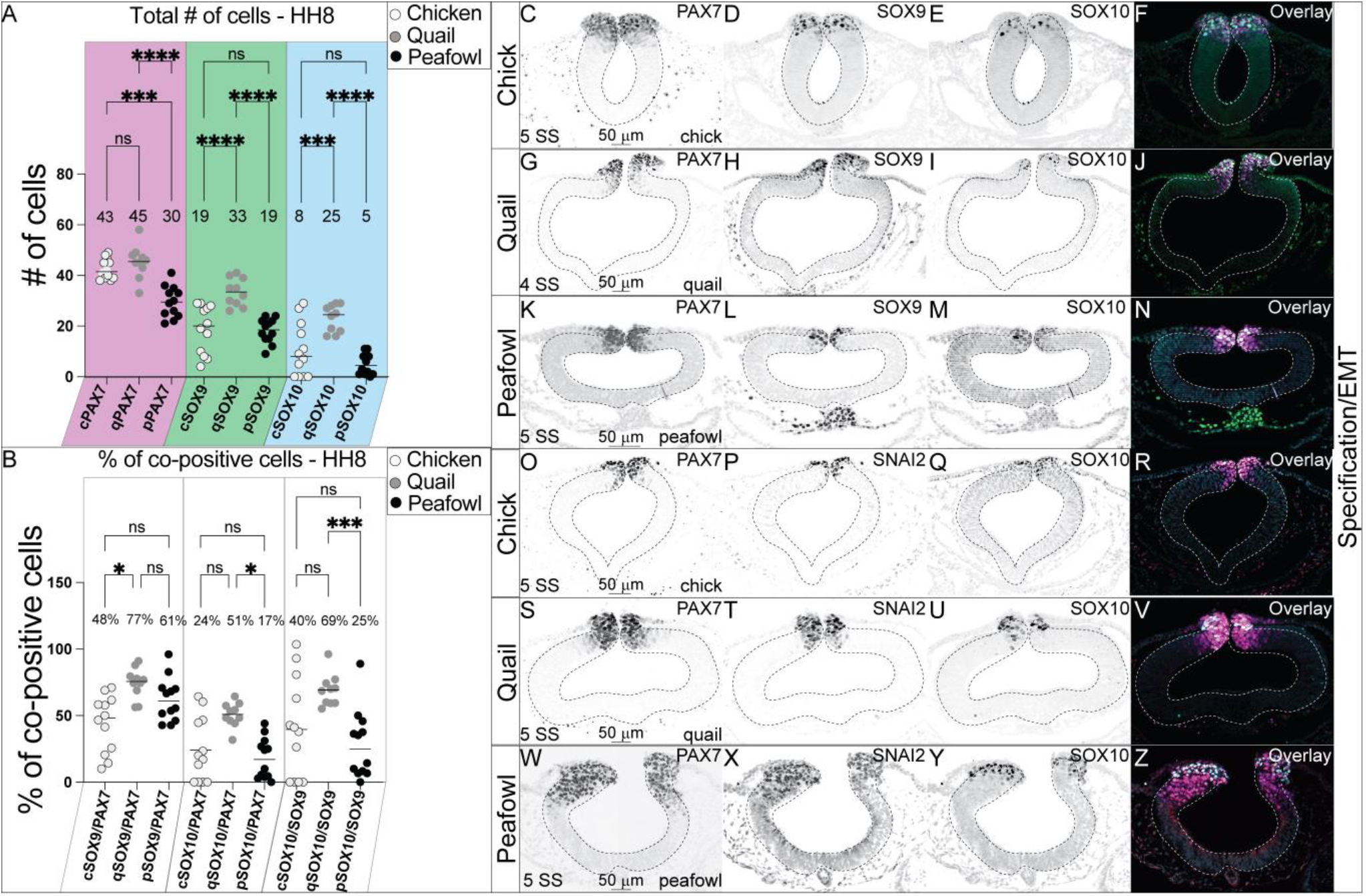
Comparative expression of PAX7, SOX9, SOX10, and SNAI2 during specification in chick, quail, and peafowl. (A) The number of PAX7+ cells is not significantly different between chicken and quail at HH8, but both chick and quail have a higher number of PAX7+ cells than peafowl (p = 0.001 between quail and peafowl, p = 0.0002 between chick and peafowl, but no statistical difference was observed between chick and quail at HH8 (p = 0.990). For SOX9+ cells, quail has more than both chick and peafowl (p = 0.001 for both). For SOX10+ cells, quail has more cells than both chick and peafowl (p = 0.001 for both). An ordinary one-way ANOVA statistical test was used. (B) Quail has a higher percentage of co-positive cells at HH8 for SOX9/PAX7 compared to chick (p = 0.034) and a higher percentage of co-positive cells for SOX10/PAX7 (p = 0.015) and SOX10/SOX9 (p = 0.0009) compared to peafowl. An ordinary one-way ANOVA statistical test was used. IHC for PAX7, SOX9, and SOX10 in (C- F) chick, (G- J) quail, and (K- N) peafowl at HH8 (5 SS). IHC for PAX7, SNAI2, and SOX10 in (O- R) chick, (S- V) quail, and (W- Z) peafowl at HH8 (5 SS). Black and white images are single protein expression at indicated stage and color images are overlay of all three markers. Scale bar is 50 μm and is indicated in first panel of each row.

**Figure 10.**
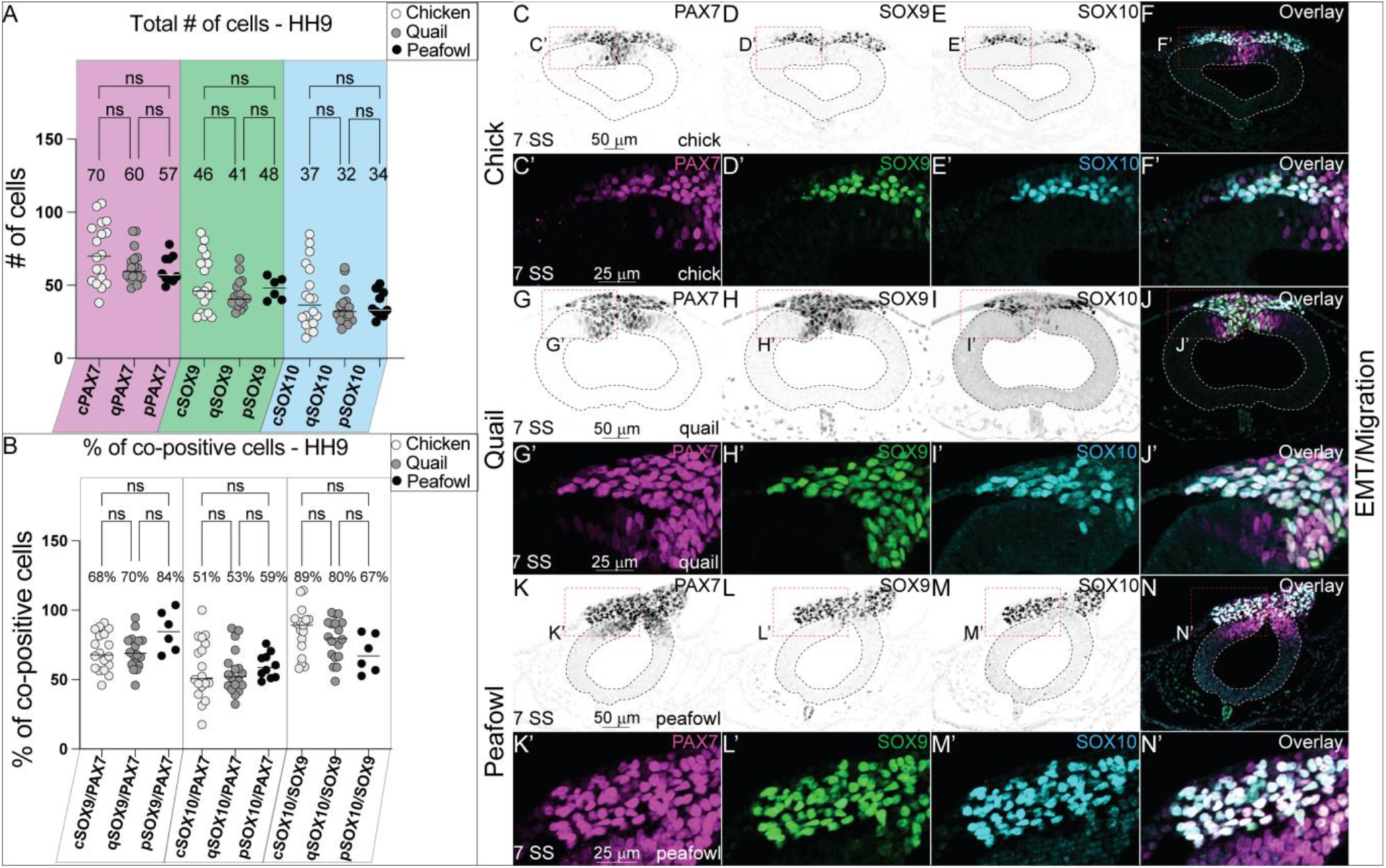
Comparative expression of PAX7, SOX9, SOX10, and SNAI2 during EMT in chick, quail, and peafowl. (A) The total number of cells for PAX7, SOX9, and SOX10 are not statistically significant between chicken, quail, and peafowl. An ordinary one-way ANOVA statistical test was used (PAX7, p = 0.675 – 0.999, SOX9, p = 0.878 – 0.999, SOX10, p = 0.895 – 0.999). (B) Percentage of co-positive cells at HH9 between chicken, quail, and peafowl shows no statistical difference. An ordinary one-way ANOVA statistical test was used (SOX9/PAX7, p = 0.9999, SOX10/PAX7, p = 0.9998, SOX10/SOX9, p = 0.8684). IHC for PAX7, SOX9, and SOX10 in (C- F’) chick, (G- J’) quail, and (K- N’) peafowl at HH9 (7 SS). Black and white images are single protein expression at indicated stage and color images are overlay of all three markers. Red dashed boxes indicate zoomed images in (C’- F’, G’- J’, and K’- N’). Neural tubes are indicated by dashed lines. Scale bar is 50 μm and is indicated in first panel of each row.

During EMT (HH9, 7-9 SS), the total number of NC cells expressing PAX7, SOX9, and SOX10 individually or together are no longer significantly different between chick, quail, or peafowl (Fig. 10A, 10C- 10N’). At this stage, the early migrating NC cell populations look very similar (Fig. 10F, 10J, 10N) in all organisms, and the percent of cells co-positive for PAX7 cells co-expressing SOX9 is 68% in chick, 70% in quail, and 84% in peafowl, but is not statistically significant (Fig. 10B, 10C, 10C’, 10D, 10D’, 10G, 10G’, 10K, 10K’, 10L, 10L’). At the same stage, the percent of PAX7+ NC cells co-expressing SOX10 is 51% in chick, 53% in quail, and 59% in peafowl and is not statistically different (Fig. 10B, 10C, 10C’, 1ED, 10E’, 10I, 10I’, 10K, 10K’, 10M, 10M’). Further, at HH9, the percent of SOX9 cells that co-express SOX10 is 89% in chick, 80% in quail, and 67% in peafowl, but the ratios are not statistically different from each other (Fig. 10B, 10D–F, 10G–J, 10L–N). Our results support the proposed hierarchical NC gene regulatory network (GRN) that begins with PAX7 expression in a large population of NC progenitors followed by the succession of the NC specifiers SOX9, SNAI2, and SOX10 (Meulemans and Bronner-Fraser, 2004; Nikitina et al., 2008; Steventon et al., 2005); however, they suggest that the timing of these proteins differs between species.

### Differential timing and expression during differentiation

Based on their differential rates of specification and migration as identified by early expression of NC specifiers and faster migration in quail (Figs. 4, 6, 8, 9), we hypothesized that there may be differences in protein localization during NC differentiation. Quail, chick, and peafowl embryos were cultured to HH15 (25 SS) and IHC for SOX9, SOX10, and PAX7 was performed (Fig. 11). We noted some anatomical differences in the developing embryos (heart size, branchial arch (BA) patterning, and anterior prominence) (Fig. 11A- 11C, 11A’- 11C’). However, true differences arose at the molecular level. Unlike chick and peafowl, quail embryos express SOX10 in the eye and BA regions (Fig. 11A’), and only weakly SOX9 in the trigeminal ganglia (TG) (Fig. 11A), while both proteins are strongly expressed in the chick and peafowl TG (Fig. 11B, 11C). Also, peafowl embryos strongly express SOX9 in their otic vesicles (OV), while the others only express SOX10 (compare Fig. 11C to 11A, 11B). These analyses suggest that at the same HH stage during differentiation phases, there are differences in avian developmental program markers at the molecular level.

**Figure 11.**
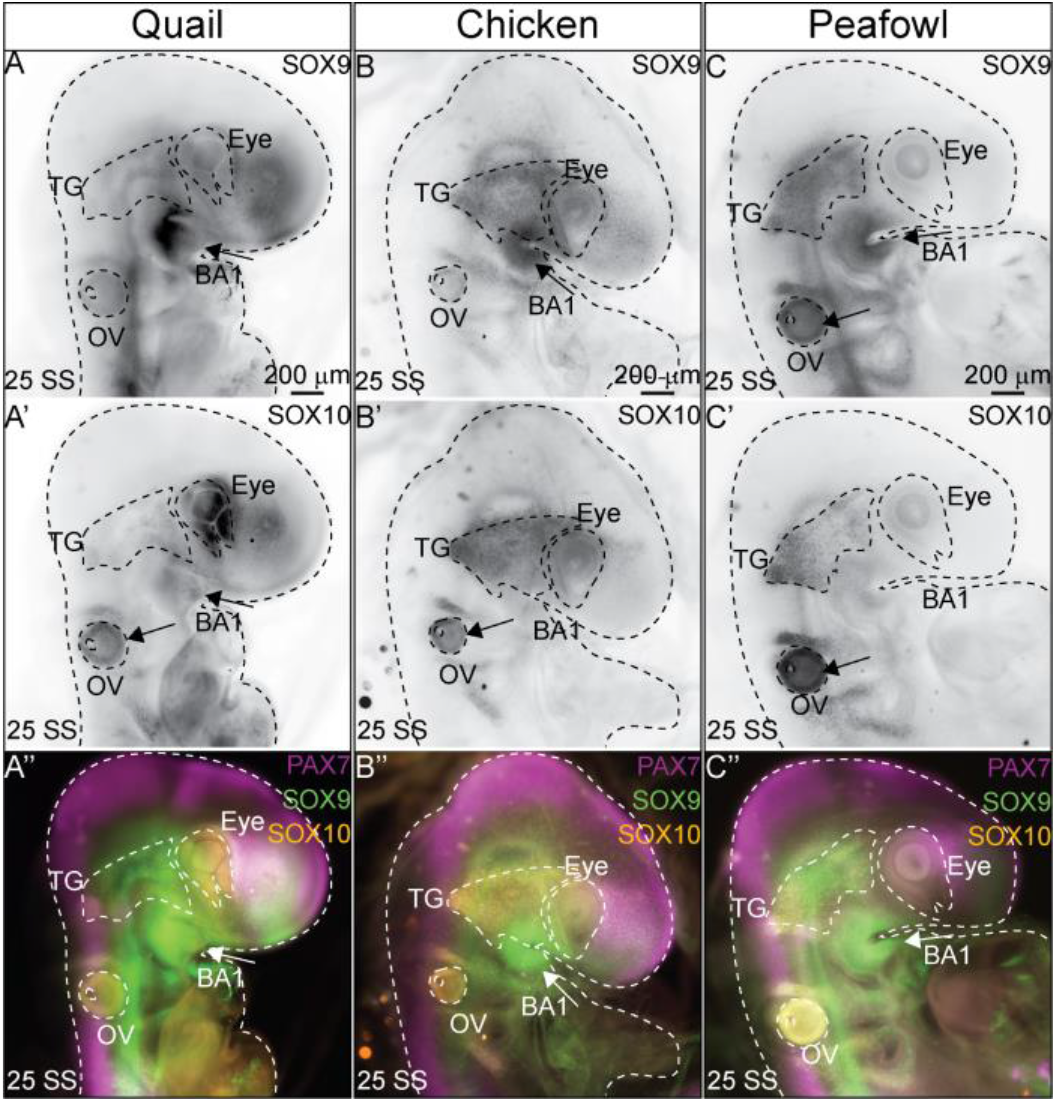
SOX9/SOX10 localization during differentiation differs between avian species. (A-C’) IHC in (A’A”) quail, (B’B”) chick, and (C-C”) peafowl embryos for (A-C) SOX9, (A’-C’) SOX10, and (A”-C”) overlay with PAX7 in 25 SS embryos. Differences: SOX9 is expressed strongly in the first branchial arch (BA) in all three organisms, but co-localizes with SOX10 only in the quail BA1. SOX10 is strongly expressed in the quail eye, but not chick and peafowl. SOX9 is prominently expressed in the peafowl otic vesicle (OV), but not quail or chick. The trigeminal ganglia (TG) strongly expresses SOX9 and SOX10 in chick and peafowl, but expression is weak in quail. Embryos are outlined with dashed lines. Scale bar is 200 μm and is indicated in top panel of each row.

## 4. DISCUSSION

We compared the spatiotemporal localization and number of PAX7, SOX9, SNAI2, and SOX10+ cells in chick, quail, and peafowl embryos from early neurula stages (HH7) to late migration and differentiation stages (HH15). We have identified that the spatial location of specified NC cells is conserved in chick, quail, and peafowl embryos. Specifically, all three organisms develop NC crest cells within the dorsal neuroepithelium, and these cells undergo a true EMT: delaminating from the neural tube, collectively migrating, and undergoing mesenchymalization during migration (Fig. 2, 4, 6, 8). In addition, although PAX7 protein expression and timing appears conserved in all species at HH7 in the neural plate border, the early onset of quail SNAI2 protein differs from expression in chick and peafowl (Fig. 3, 4) (Taneyhill et al., 2007). Further, in quail, SOX9, SNAI2, and SOX10 are expressed 1- 2 SS stages earlier than in chick and peafowl, NC cells appear to undergo EMT at an earlier stage, and the quail cells migrate further and faster. For the most part, the amino acid sequences of these proteins are highly conserved between all three species, with peafowl and quail sequences more similar to each other than those in quail (Fig. 1, 3, 5). However, the SOX10 amino acid sequence exhibits intriguing differences (Fig. 7) that we intend to study in the future. There appears to be a similar program at play that regulates specification and EMT of NC cells, but how these factors are related hierarchically in each organism remains unclear. Thus, this work provides a framework to define similarities and differences in the molecular mechanisms regulating NC cell development between closely related species with different external phenotypes and developmental timelines.

### Differences in developmental timing

While analyzing the expression of each protein, we found that quail not only expressed SNAI2, SOX9, and SOX10 at earlier equivalent developmental time points than chick and peafowl, but quail NC cells also appeared to migrate out of the neural tube earlier than chick and peafowl (HH8 versus HH9) (Fig. 2, 4, 6). This expedited NC development was particularly interesting as we identified that at HH8, quail neural tubes were significantly larger in diameter when compared to chick, but by HH9, this difference was no longer observed. In addition, we also found that at HH8, quail embryos had more SOX9 and SOX10+ cells. Taken together, we hypothesize that due to a larger neural tube and shorter developmental timeline in quail, NC GRN proteins are likely activated earlier to initiate earlier cell migration out of the neural tube. These results are consistent with the scaling pattern differences seen in the ventral neural tube between chick and zebra finch (Uygur et al., 2016).

When comparing ventral neural tube patterning between the zebra finch and the chick, the authors found that ventral neural tube patterning in the zebra finch occurs on a smaller scale and over a shorter time than in chick and that these differences in scaling are in response to a gradient of the morphogen Sonic hedgehog (Shh) (Uygur et al., 2016). Although early chick neural tube patterning (Ericson et al., 1997; Luo et al., 2006) and NC specification and EMT (Martik and Bronner, 2017; Rogers and Nie, 2018) are well characterized, less is known about the factors that drive NC induction and EMT in quail (Schneider, 2018), and virtually nothing has been previously shown about embryogenesis in general or NC-specific development in peafowl embryos.

Links between signaling molecules like Wnt, Shh, and BMP, and dorsoventral patterning, including NC formation, are well established in frog and chick (Bhattacharya et al., 2018; Chalpe et al., 2010; Garnett et al., 2012; Ossipova and Sokol, 2011; Piloto and Schilling, 2010; Shi et al., 2011). Further, cooperation between SOX9, SNAI2, and BMP signaling drives NC specification and EMT in chick and quail (Sakai et al., 2006; Zhang and Klymkowsky, 2009), however, there is a lack of information about the expression, timing, or signaling of these morphogens in other avian species during NC specification and EMT. Based on the rapid onset of quail NC specification and EMT, it is likely that the signals driving these processes also have earlier onset.

Recent work also focused on scaling in wing patterning between differently sized avian species, including quail, chick, and turkey (Stainton and Towers, 2021). The authors compared wing pattern growth rates between quail and chick and found that quail limb developmental timing is accelerated by 12 hours compared to chick. Between 0-12 hours of observation, quail wing buds grew at a significantly faster rate than chick wings and had faster proximodistal specification (Stainton and Towers, 2021). In addition, they identified that retinoic acid (RA) is responsible for species-specific developmental timing and wing scaling patterns (Stainton and Towers, 2021).

Together, these cross-species studies provide mechanisms that explain differences in developmental timing and species-specific scaling to better understand developmental growth between closely related avian species. Further studies are necessary to understand the mechanisms which drives earlier quail NC cell development compared to chick. Compared to late stage limb development, NC specification (HH8) and EMT (HH9) occur significantly earlier in development. We identified that quail NC cells were specified approximately 4-6 hours faster than chick and 8-10 hours faster than peafowl based on time of incubation. Quail embryos incubate for 17 days, whereas chick embryos incubate for 20 and peafowl for 30 days prior to hatching, so it is logical that quail undergo faster development. However, we did not identify a specific linear scale of development based on normal developmental timelines or hatched size of embryo (quail<chick<peafowl), rather, quail NC cells appeared to develop more quickly than the other avians despite their somite-based staging. Further, at the early stages analyzed, the three types of embryos were not significantly different in overall size (Figs. 1, 3, 5, 7). Our results support the idea of scaled developmental processes, but also suggest that quail NC cell development is more rapid regardless of embryonic size.

### Heterogeneous protein expression

To further quantify the similarities and differences in NC cell development between quail and chick, we counted the total number of cells, and the proportion of cells co-expressing different factors. We identified that NC cells have spatially and temporally heterogeneous NC protein expression, and that the population is not a homogenous cell population, complementing previous studies using single cell RNA-sequencing at NC specification stages (HH8) and spatial genomic analysis (SGA) at NC EMT stages (HH9) in chick embryos (Lignell et al., 2017; Williams et al., 2019). Single cell RNA-sequencing (scRNA-seq) and hybridization chain reaction (HCR) analysis performed at HH8-9 and HH10 identified high levels of *Pax7* and *Snai2* transcripts that decreased from HH8 to HH10, and that chromatin accessibility changes for NC specifiers between HH8 and HH10, suggesting that cis-regulatory landscapes are dynamic during different stages of NC development (Williams et al., 2019). SGA, a quantitative single cell profiling technique, confirmed that markers of premigratory NC cells inhabit multiple stem cell niches within the dorsal neural tube, but that these groups are spatially distinct (Lignell et al., 2017).

Our multiplexed spatiotemporal protein expression analyses support prior work demonstrating hierarchical temporal NC GRN gene expression with PAX7 protein expressed earliest in the neural plate border followed by SNAI2, then SOX9, and finally, SOX10 (Fig. 1, 3, 5, 7) (Khudyakov and Bronner-Fraser, 2009). However, not all PAX7+ cells express NC specifier proteins. Rather, it is clear that only small subsets of premigratory PAX7+ cells actively express SOX9 and SOX10 proteins during specification (HH8) for example (Fig. 9), and in quail, the co-expressing subset is significantly larger at HH8 (74% versus 44%). The early expression of bonafide NC markers, SOX9 and SNAI2, in quail versus chick and peafowl suggests that quail NC cells are specified earlier, which is supported by their early migration, but also may dictate novel protein functions and protein-protein interactions due to the different embryonic environments (Fig. 4, 6, 7, 8). Future work will focus on defining species-specific roles of these factors. These results not only highlight initial differences in the timing of protein expression of NC specifiers between avian species, but also demonstrates that gene expression studies should be complemented by protein expression analyses. Transcript expression implies a developmental state, but ultimately, the protein is the factor that drives cell state changes. Work in multiple systems has identified that mRNA and protein expression levels are not always correlated (Gry et al., 2009; Guo et al., 2008; Koussounadis et al., 2015). However, mRNA expression analyses remain strong indicators of changing developmental states for some cell types and would be bolstered by paired protein analyses.

### Sequence similarities and differences

Genome sequence analyses of multiple avian species places peafowl closest to chick in a molecular phylogeny created using homologous proteins (Dhar et al., 2019). Our work focused on identifying similarities and differences in well-studied NC transcription factors and supports these relationships. Specifically, we identified that the amino acid sequences of three out of four NC transcription factors (PAX7, SNAI2, SOX9) show that the peafowl and chick sequences are 100% similar (Figs. 1, 3, 5, Supp, Figs. 1, 2, 3). Multiple sequence alignment of DNA and amino acid sequences of the NC showed between 80%-99% similar identity between species. However, the SOX10 amino acid sequence differs in all three organisms. Strikingly, quail SOX10 has 104 additional amino acids on its N-terminus compared to chick and peafowl and 6/8 amino acid substitutions are radical replacements, where the amino acid is exchanged into another with different properties. There are multiple examples of biologically relevant single amino acid point mutations that dictate changes in protein sequence, structure, and function during disease and development (Airoldi et al., 2010). The classic example of a single amino acid change resulting in disease is that of Sickle Cell Anemia (Coleman and Inusa, 2007). Specific to neural crest cells, work in zebrafish identified that a single amino acid in the receptor tyrosine kinase protein, ERBB3, is necessary for its role in the formation of normal *Sox10*+ NC cells (Buac et al., 2008). In addition, a nonsynonymous mutation in a single amino acid in the LBH transcription factor in Cichlids is associated with alternative NC development and craniofacial adaptation (Powder et al., 2014). Therefore, although the spatial localization of PAX7, SNAI2, SOX9, and SOX10 proteins is similar between species, the differences in the timing of expression and their amino acid sequences provide a possibility of dissimilar folding, interactions, or functions in quail versus chick and peafowl, which result in differential development and dissimilar NC-related phenotypes.

## ACKNOWLEDGEMENTS

We thank our colleagues from the Rogers Lab at UC Davis in the Department of Anatomy, Physiology, and Cell Biology who provided insight and expertise. Huge thank you to Dr. Pauline Perez for providing fertilized peafowl eggs for our study. The authors declare no competing financial interests.

## AUTHOR CONTRIBUTIONS

Conceptualization: CDR and BM. Data Curation (experiments and imaging): CDR, BM, CA, and CE. Formal Analysis (cell counts, statistics, bioinformatics): BM, CDR, ACA, CA, CG. Funding Acquisition: CDR. Methodology: CDR, BM, ACA, CE. Writing: BM, CDR. Supervision: CDR.

---

**Supp. Figure 1.**
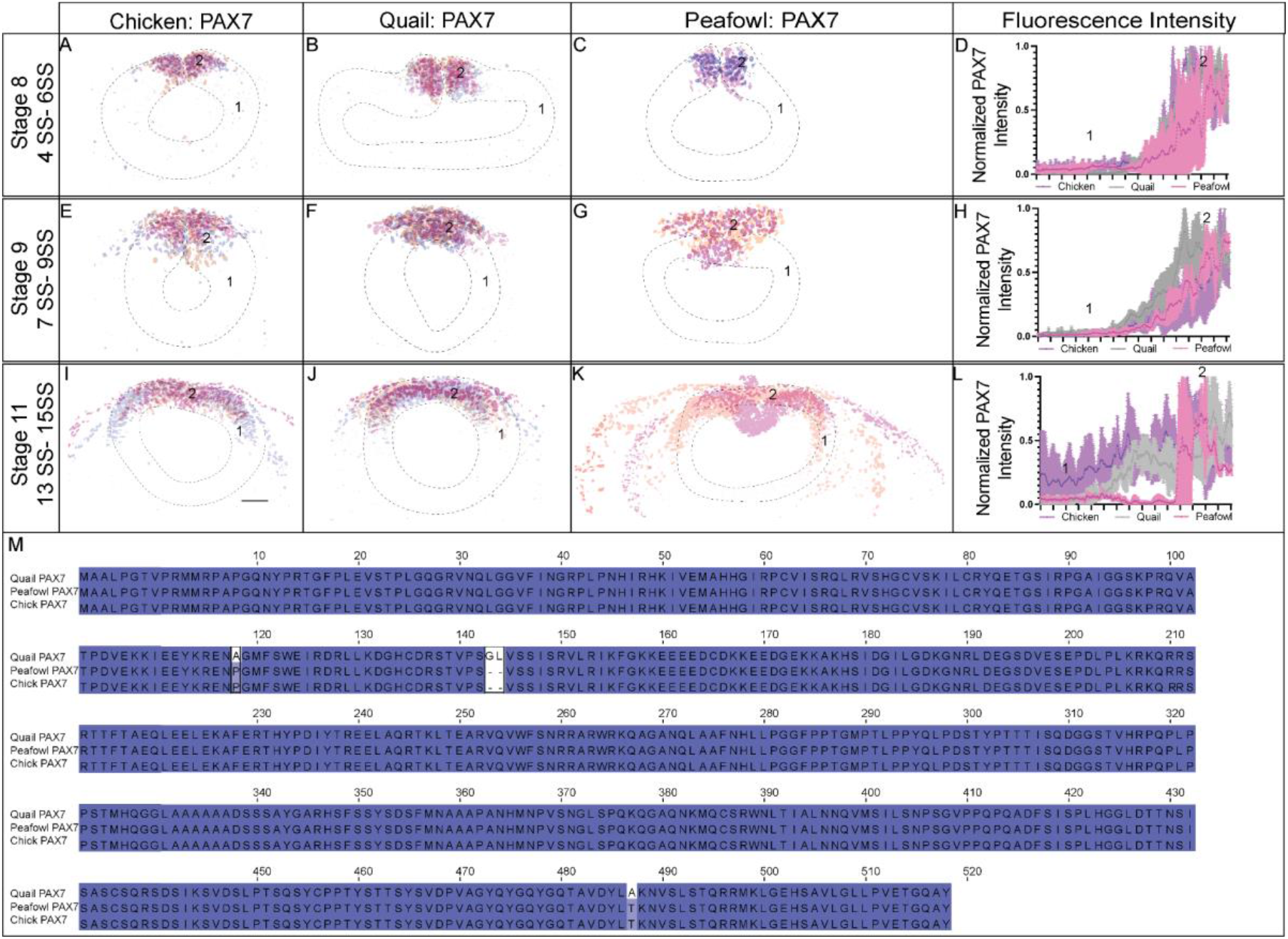
Fluorescence intensity of PAX7+ cells from medial to lateral in chick, quail, and peafowl. (A) Chicken PAX7 section overlay for HH8 (N=3). (B) Quail PAX7 section overlay for HH8 (N=3). (C) Peafowl PAX7 section overlay for HH8 (N=2). (D) PAX7 fluorescence intensity graph. Chicken (purple), quail (gray), and peafowl (pink) show lowest intensity in ventrolateral neural tube (1) (mean intensity for chicken = 0.055, 0.006 for quail, and 0.052 for peafowl p = 0.999 for all comparisons) and highest intensity in the dorsal neural tube (2) (mean intensity for chicken = 0.836, 0.787 for quail, and 0.298 for peafowl, p=0.999, 0.377, and 0.1897 for chicken to quail, quail to peafowl and chicken to peafowl, respectively). (E) Chicken PAX7 section overlay for HH9 (N=3). (E) Quail PAX7 section overlay for HH9 (N=3). (F) Peafowl PAX7 section overlay for HH9 (N=2). (H) PAX7 fluorescence intensity graph. Chicken (purple), quail (gray), peafowl (pink) show lowest intensity in ventrolateral neural tube (1) (mean intensity for chicken = 0.023, 0.037 for quail, and 0.022 for peafowl p = 0.999 for all) and highest intensity in the dorsal neural tube (2) (mean intensity for chicken = 0.377, 0.673 for quail, and 0.540 for peafowl p = 0.999 for all). (I) Chicken PAX7 section overlay for HH11 (N=3). (J) Quail PAX7 section overlay for HH11 (N=3). (K) Peafowl PAX7 section overlay for HH11 (N=2). (L) PAX7 fluorescence intensity graph. Chicken (purple) has PAX7 expression in the ventrolateral neural tube (1) (mean intensity for chicken = 0.183, 0.018 for quail, and 0.0382 for peafowl p = 0.999 for all) that continues to the dorsal neural tube (2) (mean intensity for chicken = 0.391, 0.552 for quail, and 0.291 for peafowl p = 0.9 for all). Quail (gray) has less PAX7 expression in the ventrolateral neural tube (1) and highest expression in the dorsal neural tube (2), similar to peafowl (pink). An ordinary one-way ANOVA statistical test was used to compare mean intensity values. (M) Multiple sequence alignment using quail, peafowl, and chick PAX7 amino acid sequences demonstrated that peafowl and chick sequences are 100% identical while the quail sequence has an alanine at position 118 instead of the proline and has glycine and leucine inserted at positions 143 and 144, respectively.

**Supp. Figure 2.**
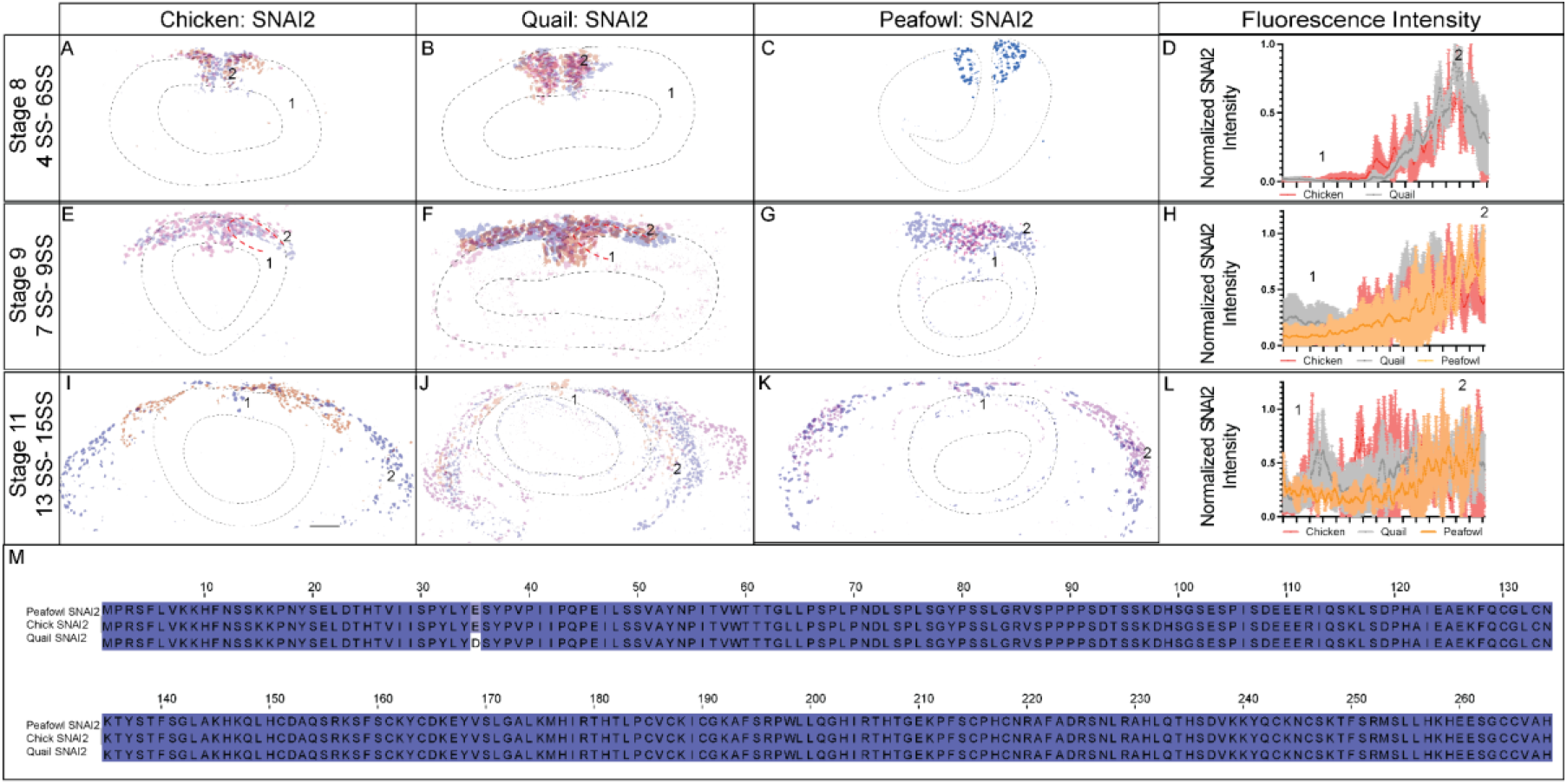
Fluorescence intensity of SNAI2+ cells from medial to lateral in chick, quail, and peafowl. (A) Chicken SNAI2 section overlay for HH8 (N=3). (B) Quail SNAI2 section overlay for HH8 (N=3). (C) Peafowl SNAI2 section for HH8 (n= 1). (D) SNAI2 fluorescence intensity graph. Chicken (red), quail (gray), and peafowl (orange) show lowest intensity in ventrolateral neural tube (1) (mean intensity for chicken = 0.0134 and 0.005 for quail, p = 0.999) and highest intensity in the dorsal neural tube (2) (mean intensity for chicken = 0.449 and 0.827 for quail, p = 0.284). (E) Chicken SNAI2 section overlay for HH9 (N=2). (F) Quail SNAI2 section overlay for HH9 (N=3). (G) Peafowl SNAI2 section overlay HH9 (N=2). Red dashed lines indicate direction of measurement from points 1 to 2. (H) SNAI2 fluorescence intensity graph. Chicken (red) and peafowl (orange) have low intensity in ventrolateral neural tube (1) (mean intensity for chicken = 0.043, 0.228 for quail, and 0.083 for peafowl p = 0.999 for all) but high expression in laterally migrating cells (2) (mean intensity for chicken = 0.372, 0.611 for quail, and 0.774 for peafowl p = 0.950 for chicken and quail, p = 0.547 for chicken and peafowl, and p = 0.000 for quail and peafowl). Quail (gray) has some SNAI2 expression in dorsal neural tube (1) with increasing expression in migrating cells (2). (I) Chicken SNAI2 section overlay for HH11 (N=2). (J) Quail SNAI2 section overlay for HH11 (N=3). (K) Peafowl SNAI2 section overlay for HH11 (N=2). (L) SNAI2 fluorescence intensity graph. Chicken (red) has SNAI2 expression in the dorsal neural tube (1) (mean intensity for chicken = 0.109, 0.194 for quail, and 0.207 for peafowl, p = 0.999 for all) that continues in laterally migrating cells (2) (mean intensity for chicken = 0.452, 0.573 for quail, and 0.679 for peafowl, p = 0.999 for all). Quail (gray) has some SNAI2 expression in the dorsal neural tube (1) but expression is highest in laterally migrating cells (2). Peafowl (orange) has little expression in dorsal neural tube (1), with higher expression in migrating cells (2). An ordinary one-way ANOVA statistical test was used to compare mean intensity values. (M) Multiple sequence alignment using peafowl, chick, and quail SNAI2 amino acid sequences demonstrated that peafowl and chick sequences are 100% identical while the quail sequence has an aspartic acid instead of a glutamic acid at position 135.

**Supp. Figure 3.**
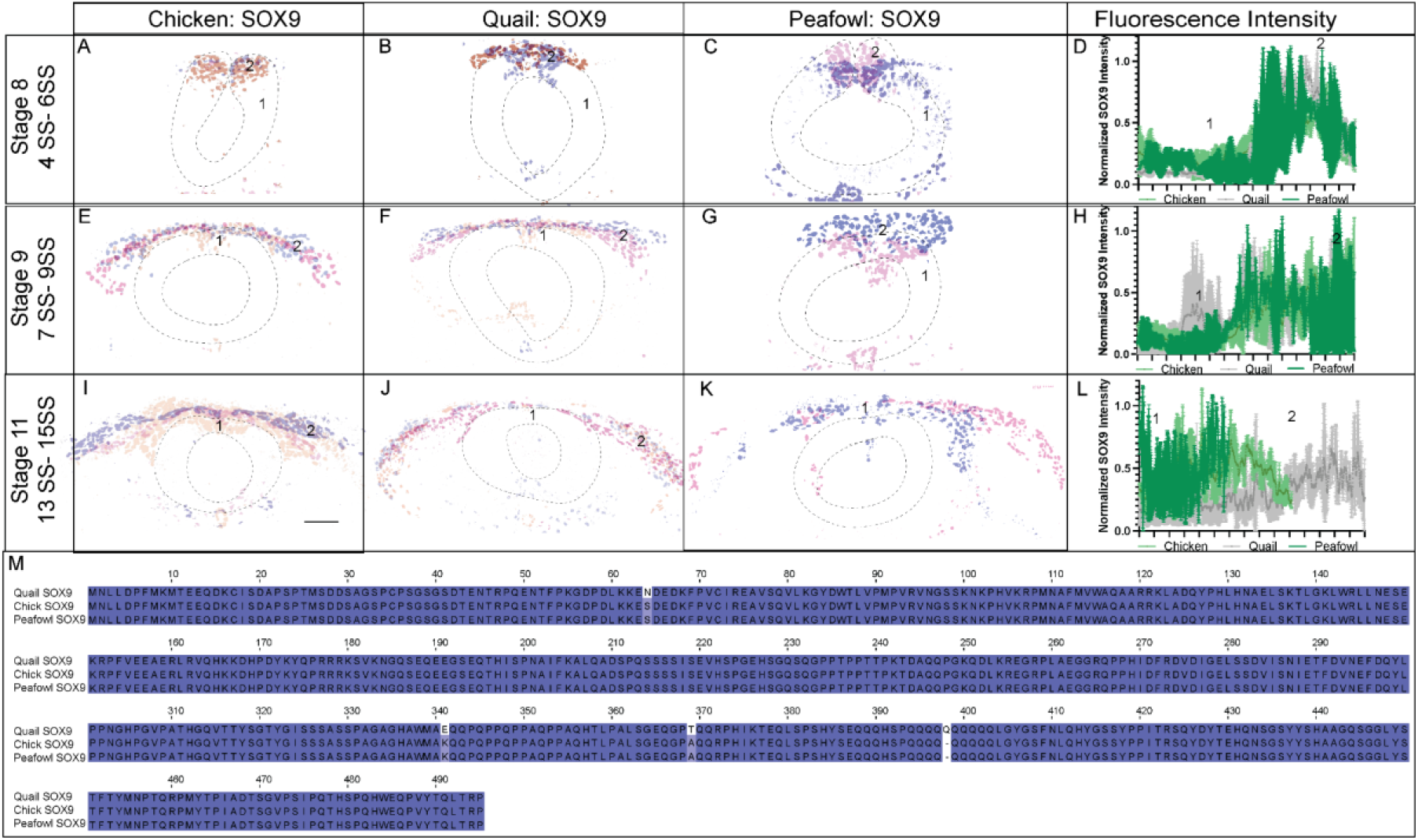
Fluorescence intensity of SOX9+ cells from medial to lateral in chick, quail and peafowl. (A) Chicken SOX9 section overlay for HH8 (N=2). (B) Quail SOX9 section overlay for HH8 (N=2). (C) Peafowl SOX9 section overlay for HH8 (N=2). (D) SOX9 fluorescence intensity graph. Chicken (green), quail (gray), and peafowl (dark green) show lowest intensity in ventrolateral neural tube (1) (mean intensity for chicken = 0.152 and 0.084 for quail, and 0.128 for peafowl, p = 0.999 for all) and highest intensity in the dorsal neural tube (2) (mean intensity for chicken = 0.513, 0.646 for quail, and 0.502 for peafowl, p = 0.999 for all). (E) Chicken SOX9 section overlay for HH9 (N=3). (F) Quail SOX9 section overlay for HH9 (N=3). (G) Peafowl SOX9 section overlay for HH9 (N=2). (H) SOX9 fluorescence intensity graph. Chicken (green) and peafowl (dark green) have low expression in dorsal neural tube (1) (mean intensity for chicken = 0.0835, 0.337 for quail, and 0.1 for peafowl, p = 0.999 for all) and continues laterally in migrating cells (2) (mean intensity for chicken = 0.451, 0.456 for quail, and 0.532 for peafowl, p = 0.999 for all). Quail (gray) has some SOX9 expression in dorsal neural tube (1) with increasing expression in migrating cells (2). (I) Chicken SOX9 section overlay for HH11 (N=3). (J) Quail SOX9 section overlay for HH11 (N=3). (K) Peafowl SOX9 section overlay for HH11 (N=2). (L) SOX9 fluorescence intensity graph. Chicken (green) has SOX9 expression in the dorsal neural tube (1) (mean intensity for chicken = 0.321, 0.167 for quail, and 0.264 for peafowl, p = 0.996 for all) that continues in laterally migrating cells (2) (mean intensity for chicken = 0.246 and 0.360 for quail, p = 0.9910). Quail (gray) has less SOX9 expression in the dorsal neural tube (1) as expression is highest in laterally migrating cells (2). Quail SOX9 fluorescence intensity expands beyond chicken and peafowl due to cells that have migrated further in quail compared to chicken. An ordinary one-way ANOVA statistical test was used to compare mean intensity values. (M) Multiple sequence alignment using peafowl, chick, and quail SOX9 amino acid sequences demonstrated that peafowl and chick sequences are 100% identical while the quail sequence has an asparagine instead of an serine at position 64, a glutamic acid instead of a lysine at position 341, a threonine instead of an alanine at position 369, and an inserted Q at position 398.

**Supp. Figure 4.**
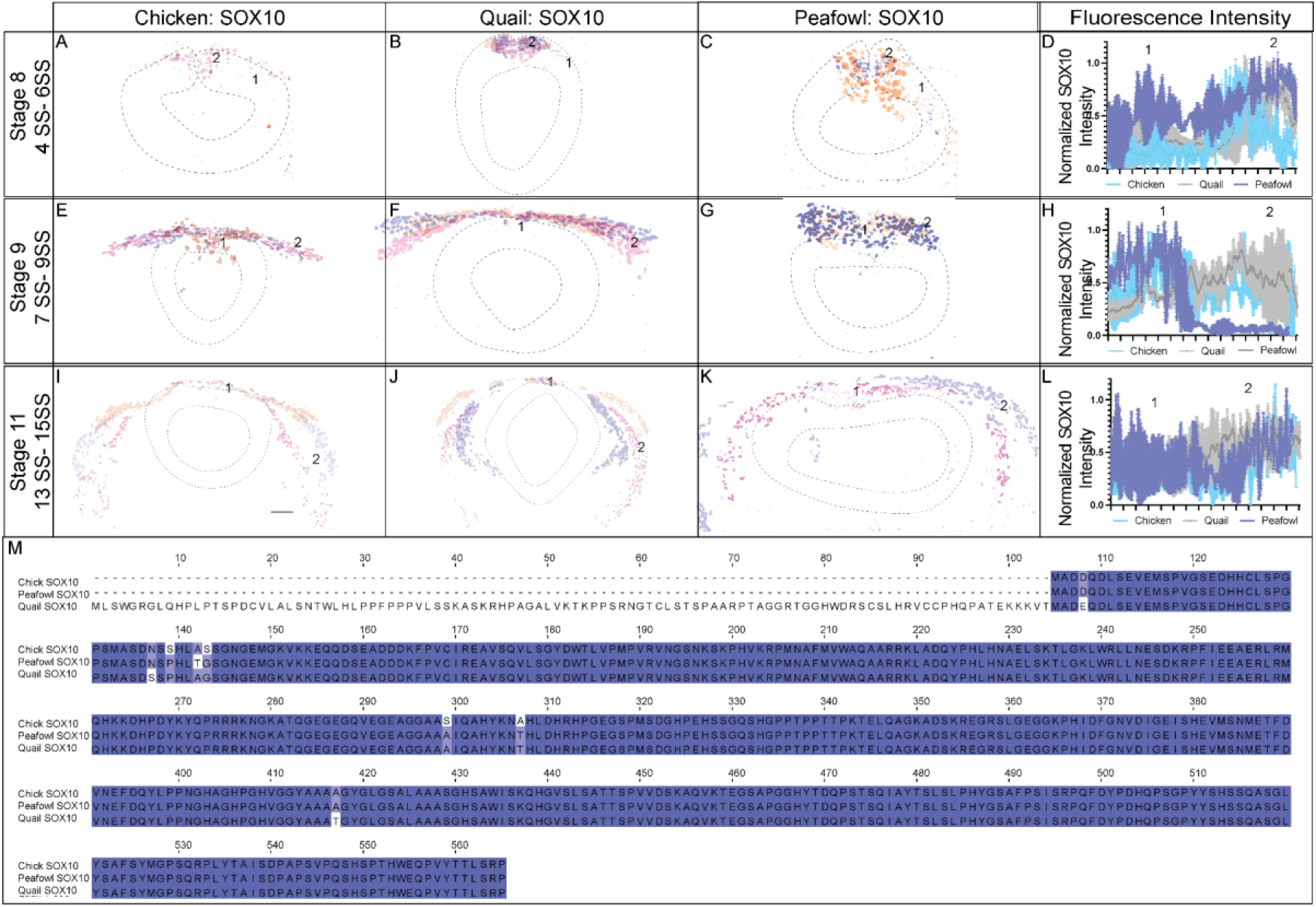
Fluorescence intensity of SOX10+ cells from medial to lateral in chick, quail, and peafowl. (A) Chicken SOX10 section overlay for HH8 (N=3). (B) Quail SOX10 section overlay for HH8 (N=3). (C) Peafowl SOX10 section overlay for HH8 (N=2). (D) SOX10 fluorescence intensity graph. Chicken (blue) and quail (gray) show lowest intensity in ventrolateral neural tube (1) (mean intensity for chicken = 0.261, 0.284 for quail, and 0.550 for peafowl, p = 0.999 for chick and quail, p = 0.813 for chick and peafowl, and 0.997 for quail and peafowl) and highest intensity in the dorsal neural tube (2) (mean intensity for chicken = 0.308, 0.665 for quail, and 0.801 for peafowl, p = 0.872 for chick and quail, p = 0.614 for chick and peafowl, and 0.999 for quail and peafowl). Peafowl (purple) has high expression in both the ventrolateral neural tube (1) and the dorsal neural tube (2). (E) Chicken SOX10 section overlay for HH9 (N=3). (F) Quail SOX10 section overlay for HH9 (N=3). (G) Peafowl SOX10 section overlay for HH9 (N=2). (H) SOX10 fluorescence intensity graph. Chicken (blue) and peafowl (purple) have high expression in dorsal neural tube (1) (mean intensity for chicken = 0.579, 0.372 for quail, and 0.793 for peafowl, p = 0.479 for chick and quail, p = 0.999 for chick and peafowl, and p = −.681 for quail and peafowl) and continues laterally in migrating cells (2) (mean intensity for chicken = 0.415, 0.491 for quail, and 0.084 for peafowl p = 0.999 for chick and quail, p = 0.970 for chick and peafowl, and p = 0.856 for quail and peafowl). Quail (gray) has some SOX10 expression in dorsal neural tube (1) with increasing expression in migrating cells (2). (I) Chicken SOX10 section overlay for HH11 (N=3). (J) Quail SOX10 section overlay for HH11 (N=3). (K) Peafowl SOX10 section overlay for HH11 (N=2). (L) SOX10 fluorescence intensity graph. Chicken (blue) has SOX10 expression in cells that have migrated out of the dorsal neural tube (1) (mean intensity for chicken = 0.202, 0.287 for quail, and 0.437 for peafowl, p = 0.999 for all) that continues in laterally migrating cells (2) (mean intensity for chicken = 0.429, 0.671 for quail, and 0.444 for peafowl, p = 0.999 for all). Quail (gray) and peafowl (purple) have SOX10 expression in the cells migrating out of the dorsal neural tube (1) and expression is highest in laterally migrating cells (2). An ordinary one-way ANOVA statistical test was used to compare mean intensity values. (M) Multiple sequence alignment using peafowl, chick, and quail SOX10 amino acid sequences demonstrated that peafowl and chick sequences are 98.92% identical, chick and quail are 80.40% similar, and quail and peafowl sequences are 81.60% similar. Both chick and peafowl have aspartic acid instead of glutamic acid at position 64, asparagine instead of serine at position 137, and alanine instead of threonine at position 417 compared to quail. Quail and peafowl share a proline instead of a serine at position 139, a glycine instead of a serine at position 143, an alanine instead of a serine at position 299, and a threonine instead of an alanine at position 307 compared to chick. Peafowl has a unique threonine instead of an alanine at position 142 compared to chick and quail.

